# Synthesis cost is a hidden driver of convergent amino acid composition in plastid ribosomal proteins

**DOI:** 10.64898/2026.08.25.747160

**Authors:** Arnav Chaudhari, Palash Sethi, Yvemirca Vilbrun, Juannan Zhou, Liming Cai

## Abstract

Protein evolution is a walk in the evolutionary space directed by mutation and selection. While functional and structural constraints serve as the main determinant of amino acid substitution in most proteins, synthesis cost and mutational bias can also alter the direction and rate of amino acid evolution, especially in systems experiencing relaxed selection. Here, we focused on the highly expressed plastid ribosomal proteins (PRP), which comprise 58 conserved proteins encoded by both plastid and nuclear genomes. Relaxed selection has been repeatedly identified in three distantly related plant lineages, providing a valuable comparative framework to investigate the significance of synthesis cost and mutation. We first demonstrated that the hemiparasitic tribe Cymbarieae (Orobanchaceae) represented a new case where concerted cyto-nuclear rate elevation occurs in their PRP. Further investigation revealed convergent shifts in amino acid composition in all four plant lineages attributable to arginine-to-lysine and methionine-to-isoleucine/valine/leucine substitutions. The replacement residues were biophysically similar but had lower molecular weight and shorter side chains, which significantly destabilized protein folding as demonstrated by protein structure modeling. We found that the composition shifts ran counter to the expectation of mutational bias but were consistent with the expectation of synthesis cost minimization, which is potentially adaptive for highly expressed PRP. Further, cost minimization significantly influenced all conservative substitutions between biophysically similar amino acids but was absent in non-conservative substitutions. We thus propose cost minimization as a secondary selective drive for protein evolution in PRP, unmasked in lineages and sites with relaxed selection on their function.

## INTRODUCTION

The molecular evolution of protein sequences is influenced by a suite of selective and mutational forces. A widely accepted principle is that the rate and direction of amino acid substitution are primarily determined by the stringency of structural and functional constraints (Kimura 1968; Ohta 1973; Nei 1987). For example, transmembrane proteins and large interacting complexes have lower substitution rates because they have more stringent structural requirements compared to globular proteins or small interacting protein complexes (Tourasse and Li 2000; Fraser et al. 2002; Bloom and Adami 2003). Beyond structural constraint, mutational bias and biosynthesis cost are also frequently explored molecular drivers of protein evolution, whose effects can be quantified by amino acid composition (Graur 1985; Seligmann 2003). If mutation dominates protein evolution, the amino acid composition is expected to drift towards the background mutational bias; similarly, minimization of synthesis cost may lead to the preferred use of less costly amino acids (Singer and Hickey 2000; Barton et al. 2010). However, the effects of mutational and cost constraints are often obscured by stronger structural constraints and are therefore most evident under extreme conditions or when structural constraints are explicitly controlled (Seligmann 2003). For example, synthesis cost is negatively correlated with amino acid abundance across archaea, bacteria, and eukarya (Seligmann 2003; Swire 2007), but this relationship is strongest in nutrient-limited environments where resource conservation is under stronger selection (Mende et al. 2017). Likewise, cost minimization is stronger at sites where alternative amino acids have minimal structural consequences (Seligmann 2003) and among biophysically similar amino acid pairs in species with large effective population sizes (McShea et al. 2025). A second challenge in evaluating the role of synthesis cost is measuring the cost itself. While much work has utilized chemical energy (Craig and Weber 1998; Akashi and Gojobori 2002; Wagner 2005; Barton et al. 2010) or molecular weight (Seligmann 2003) to measure costs, they may not represent the full scope of biosynthesis investment in different cellular and environmental conditions (Barton et al. 2010). Recently, Chen and Nielsen (2022) used the biomass of enzymes required per synthesis (i.e., the protein cost) and found it outperformed chemical energy measurements in predicting amino acid abundance in yeast. Interpretations of cost-driven patterns in protein evolution thus must be made with careful consideration of how biosynthetic cost is defined.

The plastid ribosome proteins (PRP) represent an unusual opportunity to investigate the complex interplay of structural constraint, biosynthesis cost, and mutational biases. The plastid ribosome is a plant-specific organellar translation machinery consisting of approximately 58 protein subunits and 4 ribosomal RNAs (Yamaguchi et al. 2000; Yamaguchi and Subramanian 2000). Its composition and general mode of function are highly conserved across plants and even very similar to the eubacterial 70S ribosome (Subramanian 1993). The structure is further constrained by the numerous contact residues of plastid-and nuclear-encoded subunits that are under strong purifying selection for cytonuclear coevolution (Sloan et al. 2014; Sloan et al. 2018). Meanwhile, plastid ribosomes are responsible for synthesizing huge amounts of biomass in plastids, including the most abundant protein on Earth, RuBisCO (Yamaguchi and Subramanian 2000). They are thus under strong selective pressure to be produced rapidly and cost-effectively, such that ribosomal proteins have been the gold standard to identify optimal codons in a given organism (Tiller and Bock 2014; Hanson and Coller 2018). As a result, the evolution of PRP is jointly shaped by outstanding structural and synthesis cost constraints. Yet relaxed selection and high rates of non-synonymous substitution have been independently identified in *Silene* (Caryophyllaceae), Geraniaceae, the 50-kb inversion Papilionoideae (Fabaceae), and potentially other lineages (Sloan et al. 2014; Weng et al. 2016; Yang et al. 2023; Tressel et al. 2025), offering a comparative framework to track protein evolution upon lifted structural constraint. Specifically, relaxed structural constraint enables us to untangle the otherwise masked effects of synthesis cost and mutational bias.

Here, we focused on the amino acid composition and protein structural stability in plant lineages that experience relaxed selection in their PRP. We hypothesized that relaxation of structural constraint would unmask the effects of secondary selective pressures such as biosynthesis cost and even the background mutational bias. We first demonstrated that the parasitic plant tribe Cymbarieae (Orobanchaceae) represented a new plant lineage showing relaxed selection in their PRP. Cymbarieae is a member of the parasitic Orobanchaceae family and holds a critical position for understanding the transition from free-living to a heterotrophic lifestyle in plants (Cai 2023; Ma et al. 2024). It contains 13 species from 5 genera in China, Mongolia, Kazakhstan, Ukraine, Turkey, and the United States. Our previous work on organellar evolution in Orobanchaceae revealed unusually elevated evolution rates of PRP in a single member of Cymbarieae, *Monochasma sheareri* (Maxim.) Franch. ex Maxim (Cai et al. 2026). Here, we expanded taxon and nuclear gene sampling to formally test this hypothesis in tribe Cymbarieae. We then examined shifts in amino acid composition in all four plant lineages with relaxed selection of PRP. We found convergent decrease in usage of arginine and methionine caused by the replacement of biophysically similar but biosynthetically cheaper amino acids. Finally, based on the high-resolution cryo-EM structure of PRP and deep-learning-powered protein structure prediction tools (Sharma et al. 2007; Li et al. 2020; Jumper et al. 2021; Blaabjerg et al. 2023; McBride et al. 2023), we demonstrated negative structural impacts of these amino acid replacements and their significant cost-minimization effect compared to the background.

## RESULTS

### The curated Orobanchaceae PRP sequences

We curated plastid and nuclear sequences of PRP and other reference genes in Orobanchaceae. All sequences used in this study were available in Data S1. The plastid dataset contained 80 protein-coding genes from 24 representative species in Orobanchaceae (Fig. 1; Table S1). The taxon sampling spanned diverse lifestyle strategies in this parasitic family and all major tribes sensu Fischer (2004), including four of the five recognized genera within Cymbarieae: *Cymbaria* L., *Monochasma* Maxim., *Siphonostegia* Benth, and *Schwalbea* L (Fig. S1). The 80 plastid genes comprised 21 plastid-encoded PRP genes (CpPRP) and 20 plastid-encoded photosynthesis genes (CpPS), along with additional genes representing other major plastid functional categories. Loss of plastid genes has been well documented in Orobanchaceae and more recently in tribe Cymbarieae due to the evolution of parasitism (Ma et al. 2024). We similarly found prevalent gene loss and pseudogenization in non-photosynthetic holoparasites such as *Orobanche*, *Aeginetia*, and *Lathraea*, especially in their Photosystem I and II, ATP synthase, NDH complex, and RNA polymerase (Fig. 1A). In tribes Cymbarieae and Pedicularideae, frequent pseudogenization of the NDH complex and photosystem genes was identified (Fig. 1A).

**Figure 1.**
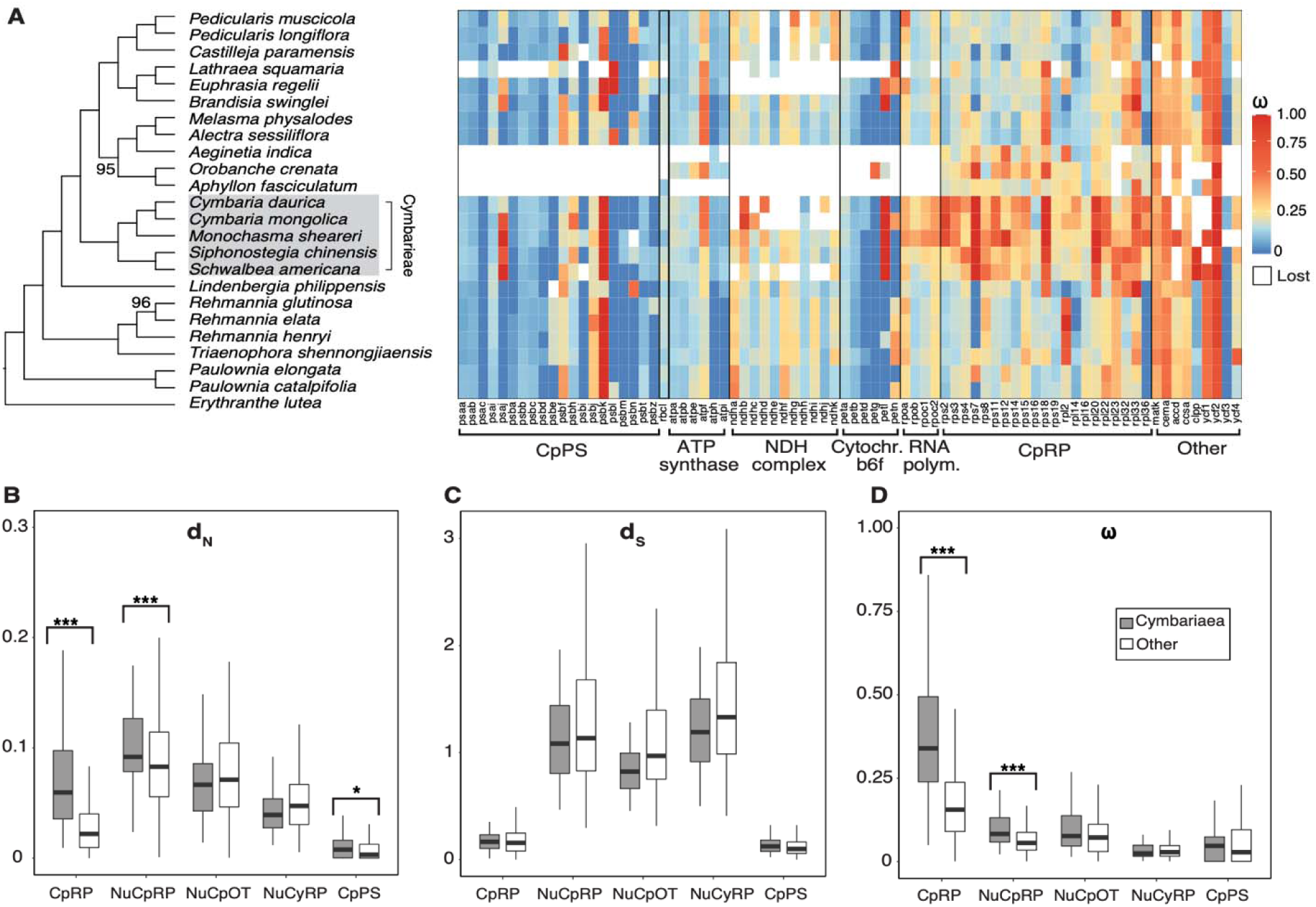
Nucleotide substitution rates in plastid-and nuclear-encoded genes across Orobanchaceae. (A) Heatmap of ω (d_N_/d_S_) values for all plastid-encoded protein-coding genes showing gene-and lineage-specific rate variation. Warmer colors indicate elevated ω values and are concentrated in the plastid-encoded ribosomal protein genes (CpPRP) of Cymbarieae. The maximum likelihood phylogeny on the left is reconstructed from concatenated plastid genes in IQTREE. Node labels indicate branch support measured by 1,000 ultrafast bootstrap and only values <100 are shown. (B–D) Boxplots of the nonsynonymous (d_N_), synonymous (d_S_), and ω values between Cymbarieae and other Orobanchaceae across five functional groups: plastid-encoded ribosomal proteins (CpPRP), plastid-encoded photosynthetic genes (CpPS), nuclear-encoded plastid-targeted ribosomal proteins (NuPRP), nuclear-encoded plastid-targeted non-ribosomal proteins (NuCpOT), and nuclear-encoded cytosolic ribosomal proteins (NuCyRP). Asterisks above boxplots indicate significance levels from Wilcoxon rank-sum tests comparing Cymbarieae to other Orobanchaceae lineages (* FDR-adjusted p-value < 0.05; ** p < 0.01; *** p < 0.001).

The nuclear dataset included published genomes and transcriptomes from 22 Orobanchaceae species that aligned with the genus-level sampling of the plastid dataset (Table S2). These genomes and transcriptomes demonstrated overall high quality and completeness with an average of 10.7% missing land plant BUSCOs (Table S2). The highest level of missing BUSCOs was found in non-photosynthetic parasites such as *Aeginetia*, *Conopholis*, and *Aphyllon* as expected because they have lost photosynthetic-related genes that are overrepresented in plant BUSCOs (Cai et al. 2021; Cai et al. 2025). We identified orthologs of 61 nuclear-encoded genes variously targeted at plastid or cytosol functions (Table S3), including 29 nuclear-encoded PRP (NuPRP), 17 nuclear-encoded cytosol-targeted ribosomal genes (NuCyRP), and 15 other nuclear-encoded plastid-targeted genes that are not involved in ribosomes (NuCpOT).

### PRP genes show elevated d_N_ and ω in Cymbarieae

To characterize selection and overall substitution rate, nonsynonymous substitution rates (d_N_) and synonymous substitution rates (d_S_) values for each gene and species were estimated using pairwise comparison with the outgroup Phrymaceae in CODEML (Fig. 1) (Yang 2007). Prior to CODEML evaluation, we bioinformatically predicted and masked C-to-U RNA editing sites in plastid genes using PREPACT v3 (Table S4) (Lenz et al. 2018). We also annotated and trimmed N-terminal peptides for nuclear genes (Table S3). Substitution rates of examined genes exhibited a significant gene-and lineage-specific pattern (Fig. 1A). Within the plastid genome, photosynthetic genes (CpPS) generally showed lower ω (median = 0.0913) compared with CpPRP (median = 0.1390)(Table S5). Within CpPRP genes, the lineage-specific shifts in ω values were most significant in Cymbarieae. Here, the median ω of CpPRP in Cymbarieae was 0.3537, which was significantly higher than the other Orobanchaceae species (median = 0.1552; Wilcoxon test FDR adjusted p-value = 4.61e^-26^), including the holoparasites (Fig. 1D). Further investigation revealed that such difference in ω was driven by elevated d_N_ values (Wilcoxon test FDR adjusted p-value = 4.77e^-19^; Fig. 1B) while the d_S_ did not differ from the other Orobanchaceae (Wilcoxon test FDR adjusted p-value = 0.58; Fig. 1C). Significantly elevated d_N_, but not ω, was also found in the CpPS of Cymbarieae relative to other photosynthetic species (Wilcoxon test FDR adjusted p-value = 0.0445). For the nuclear genome, we similarly identified significantly increased d_N_ (Wilcoxon test FDR adjusted p-value = 8.25e⁻⁴) and ω (Wilcoxon test FDR adjusted p-value = 8.25e⁻⁴) in NuPRP of Cymbarieae compared to other lineages, but not in their d_S_ (Wilcoxon test FDR adjusted p-value = 0.585). No significant rate differences were detected for NuCyRP or NuCpOT.

### Relaxed selection in Cymbarieae PRP genes

As a second measurement of selective shifts, the RELAX analysis from HyPhy v4.5 (Wertheim et al. 2015; Kosakovsky Pond et al. 2020) was applied to both individual genes and concatenated sequences grouped by function (Tables S3, S6). When grouping plastid genes by function to reduce noise (Table S6), we found that tribe Cymbarieae experienced relaxation in the ribosomal small subunit, ribosomal large subunit, Photosystem I, ATP synthase, NAD(P)H dehydrogenase, and RNA polymerase. Within the two subunits of ribosomes, 58% (n=7) of the genes in the small subunit were under significant relaxation and the concatenated sequence showed the strongest signal of relaxed selection (*k* = 1.043e^-7^; p-value = 0) compared to all protein complexes examined in our study. Similarly, 56% (n=5) of the genes in the large subunit were under relaxation. Beyond PRP, relaxation was also significant in *rbcL*(*k* = 0.678; p-value = 6.17e^-4^) and *ccsA*(*k* = 0.557; p-value = 0.011), which encode RuBisCO and Cytochrome c synthesis A, respectively.

The NuPRP genes largely mirrored this pattern in Cymbarieae (Table S3). Among the three different functional groups (NuPRP, NuCyRP, and NuCpOT), 14 out of 15 cases of relaxed selection were observed within NuPRP, which made up 48% (n=14) of its total members. These relax-selected NuPRP genes included not only structural subunits such as *RPL* and *RPS* but also ribosome binding factors such as *PSRP-1* and *PSRP-2* (Table S3).

### Convergent shifts of amino acid composition in all four plant lineages

To explore convergent trends of molecular evolution in plant lineages experiencing relaxed selection in their PRP, we expanded taxon sampling to include the 50-kb inversion clade of Papilionoideae (Fabaceae), Geraniaceae, and *Silene* (Caryophyllaceae), where concerted rate acceleration of PRP was independently identified (Table S7) (Sloan et al. 2014; Weng et al. 2016; Tressel et al. 2025). Our initial attempt to identify convergent site substitutions in these four focal clades using CSUBST (Fukushima and Pollock 2023) yielded negative results (Note S1). Instead, we discovered surprising convergent shifts in the overall amino acid composition (Fig. 2). Such convergent shifts in amino acid composition were unique to CpPRP and absent in NuPRP (Fig. S2).

**Figure 2.**
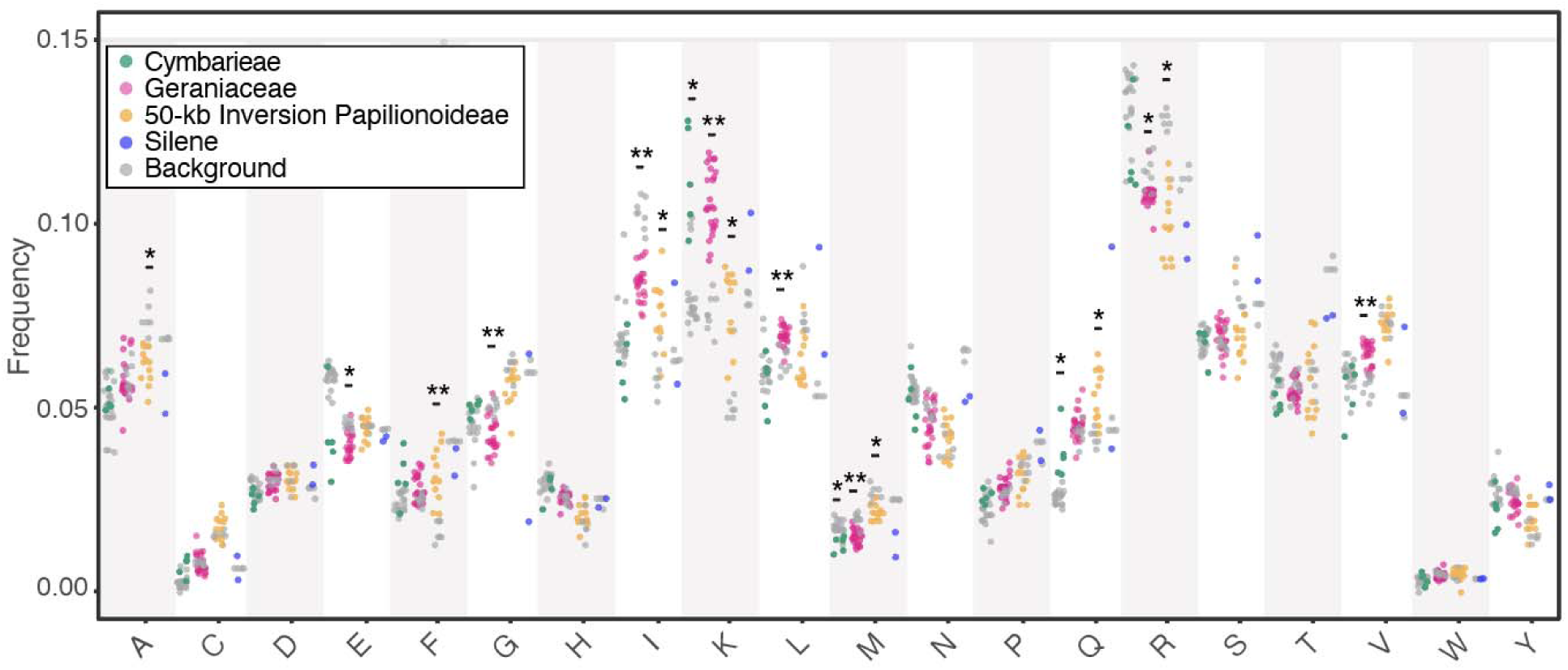
Convergent shifts in amino acid usage after relaxed selection in plastid-encoded ribosomal proteins. Frequencies of amino acid usage are contrasted between species experiencing relaxed selection (colored dots) and background reference species within the same clade (gray dots) in four plant lineages. Asterisks indicate significant differences in amino acid frequency based on Wilcoxon rank-sum tests (* FDR-adjusted p-value < 0.05; ** p-value < 0.01). Consistent increases in lysine (K) and decreases in methionine (M) are observed in all four lineages.

In CpPRP, increased use of lysine (K) was seen in all four focal lineages (Fig. 2), three of which were significant (Wilcoxon test FDR adjusted p-value <0.05). The lack of statistical significance in *Silene* was caused by its small sample size (n = 5), making none of the 20 amino acid comparisons statistically significant (Table S8). Similarly, decreased use of methionine (M) and arginine (R) was seen in all four lineages; three were significant for M and two were significant for R (Fig. 2; Table S8). Other shifts in amino acid compositions were less consistent across all taxa, including the significantly increased use of glutamine in Cymbarieae and the 50-kb inversion Papilionoideae, the reduced use of alanine in the 50-kb inversion Papilionoideae, among others (Fig. 2). We performed the same test for NuPRP, bu only 2 out of 80 comparisons showed significant differences and the direction of the shifts was inconsistent among lineages (Fig. S2).

To identify substitutions contributing to the overall shifts in amino acid composition, we used the empirical Bayesian method implemented in IQ-TREE v3.0.1 (Nguyen et al. 2015; Wong et al. 2025) to infer ancestral sequences and found prevalent R⍰K substitutions (Fig. 3). In all four lineages, REK substitution was the most abundant amino acid substitution observed in lineages experiencing relaxed selection, accounting for 3.3–8.3% of the total substitutions. Taking Cymbarieae as an example, there were 36 REK substitutions contributing to the 87 total gains of K across all internal and terminal branches. These gains outnumbered the 47 losses of K and led to the significant net increase of K compared to other reference species. In addition to R⍰K substitutions, asparagine-to-lysine (N⍰K) and glutamate-to-lysine (E⍰K) substitutions frequently ranked among the five most frequent substitutions across four plant lineages (Fig. 3), further contributing to the consistent increase of K. On the other hand, the consistent decrease of M often involved the replacement of M with isoleucine (I), leucine (L), and valine (V), and sometimes, threonine (T). Across all four lineages, there were 8.0–14.5% more losses of M than gains. Unlike R or K, substitutions involving M often only involved a few biophysically similar amino acids (Fig. 3). We found that some sites were particularly prone to these substitutions such that convergent substitutions were identified in multiple lineages. For example, in rps2, the mutation M14I/L evolved three times in Geraniaceae and Papilionoideae; and R25K also evolved three times in Cymbarieae and Geraniaceae (Fig. 4E).

**Figure 3.**
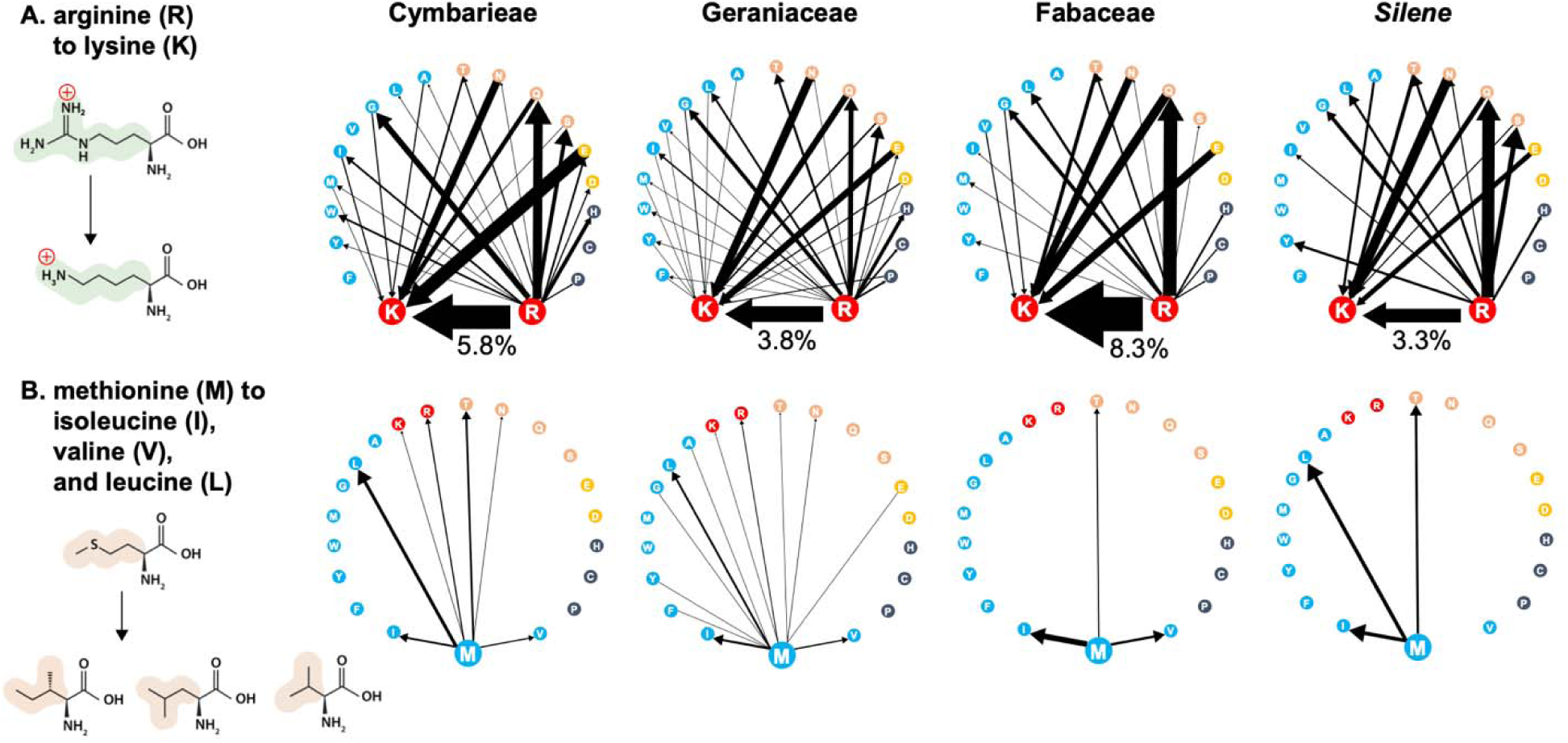
Net substitutions between pairs of biophysically similar amino acids in plastid-encoded ribosomal proteins with relaxed selection. The substitution flux is summarized by adding up reconstructed substitutions across all internal and terminal branches of lineages experiencing relaxed selection in their plastid ribosomal proteins. Arrows represent the direction of substitutions. Edge weights represent the substitution frequency normalized by the total number of substitutions in each lineage, as shown as percentages for all arginine-to-lysine substitutions. Amino acids are colored based on shared chemical properties. (A) Frequency of substitution from arginine (R) and to lysine (K) in four plant lineages. (B) Frequency of substitution from methionine (M) to isoleucine (I), valine (V), and leucine (L) in four plant lineages.

**Figure 4.**
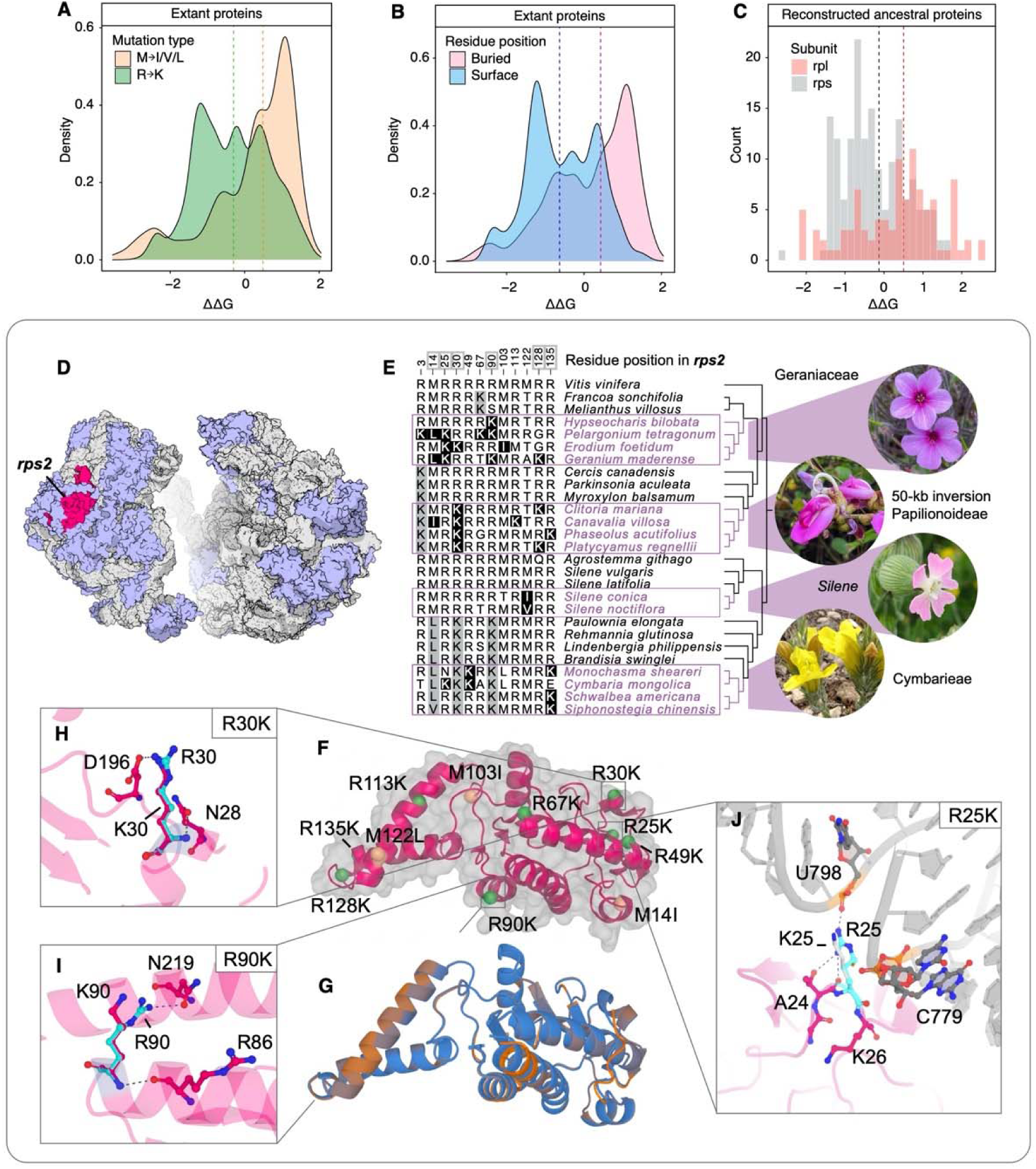
Replacement of arginine (R) and methionine (M) leads to generally destabilized structure of the plastid ribosome and a case example in *rps2*. (A-C) Impact of M⍰I/V/L and R⍰K mutations on plastid ribosome protein structure measured by free energy of protein folding (ΔΔG) in ELASPIC2. Positive ΔΔG suggests destabilized proteins. Dotted lines indicate the median value of ΔΔG for each category. (A) M⍰I/V/L mutations have stronger negative effects on protein stability compared to R⍰K mutations. ΔΔG was inferred from pairwise comparisons of protein structure from extant species with and without relaxed selection. (B) Mutations in buried residues have stronger destabilizing effects compared to surface residues. The classification of surface versus buried residues was based on relative solvent accessibility estimated by FreeSASA. (C) Mutations in the large subunit have stronger destabilizing effects than the small subunit. ΔΔG was inferred by introducing individual M⍰I/V/L or R⍰K mutations to the reconstructed ancestral protein of each lineage. (D-J) Negative structural impacts of M⍰LI/V/L and R⍰K mutations in *rps2*. (D) Cryo-EM structure of the small and large subunit of plastid ribosome from the RCSB Protein Data Bank (https://www.rcsb.org/). *rps2* is highlighted in magenta. Other ribosome proteins are shown in light blue and RNA in gray. (E) Amino acid alignment of *rps2* showing parallel M⍰I/V/L and R⍰K substitutions. Numbers on the top represent residue position and circled numbers indicate parallel substitutions in multiple lineages. Black boxes in the alignment highlight M⍰I/V/L and R⍰K substitutions in the focal lineages; gray boxes indicate ancestral M⍰I/V/L and R⍰K substitutions. The species tree on the right was subsampled from the original maximum likelihood phylogeny of concatenated plastid genes to facilitate visualization. For the full tree, see Fig. S6. (F) Protein model of *rps2* with highlighted positions of M⍰I/V/L and R⍰K substitutions in the four plant lineages. (G) Alignment of protein structure between ancestral *Silene rps2* (blue) and derived *Silene noctiflora rps2* (orange). The color gradient from blue to orange shows low to high structural difference. Given the 31 substitutions in *Silene noctiflora* compared to the reconstructed ancestral sequence of *Silene*, *rps2* is relatively robust to mutations. (H) The R30K substitution in Papilionoideae (Fabaceae) breaks the hydrogen bond with D196, thus destabilizing *rps2.* (I) The buried R90K substitution in Geraniaceae breaks the hydrogen bond with N219, thus destabilizing *rps2.* (J) The surface R25K substitution in Papilionoideae and Cymbarieae (Orobanchaceae) breaks the hydrogen bond with A24 or even 16S rRNA depending on the rotamer, thus destabilizing protein-protein or protein-rRNA interactions in the small subunit complex.

### Replacement of ancestral amino acids leads to reduced protein folding stability

To test the structural impacts of amino acid mutations such as R⍰K, we applied the deep-learning-powered ELASPIC2 to predict protein folding stability (Strokach et al. 2021). We performed two complementary sets of analyses to examine protein folding stability of 1) proteins from extant species pairs (e.g., a Cymbarieae species and its closely related reference species) and 2) reconstructed ancestral proteins with introduced mutations (e.g., reconstructed ancestral protein of Cymbarieae with introduced R⍰K mutations). Both experiments suggested that lineages marked with relaxed selection had overall destabilized protein structures. The destabilizing effect was stronger in the large subunit than in the small subunit, more pronounced for M⍰I/V/L than for R⍰K mutations, and greater in buried than in surface residues (Fig. 4A-C; Fig S3).

First, the direct comparison of protein sequences between extant species with and without relaxed selection revealed that relaxed selection was associated with increased free energy of protein folding (ΔΔG > 0; Wilcoxon test FDR adjusted p-value < 4.33e^-5^), which indicated destabilized protein structures. The large subunit was slightly more sensitive to mutation compared to the small subunit (Wilcoxon test FDR adjusted p-value = 0.53), where the median ΔΔG was 1.327 in the large subunit compared to 0.534 in the small subunit (Fig. S3A). When focusing on the effects of individual mutations, we found that M⍰I/V/L mutations were significantly more likely to be destabilizing compared to R⍰K (Fig. 4A; Wilcoxon test FDR adjusted p-value < 2.2e^-16^), where the median ΔΔG was 0.491 for M⍰I/V/L mutations and 69.1% of such mutations were predicted to destabilize the protein (ΔΔG > 0). In contrast, R⍰K mutations were more likely to be neutral or even beneficial in terms of protein stability, where the median ΔΔG was −0.299 and only 38.7% were predicted to destabilize the protein (Fig. 4A). The position of the amino acid residue also had a significant influence (Fig. 4B; Wilcoxon test FDR adjusted p-value < 2.2e^-16^), where buried residues were more sensitive to mutations (median ΔΔG = 0.431) compared to surface residues (median ΔΔG = −0.630). This at least partially explained the contrasting results between M⍰I/V/L versus R⍰K mutations because 90.8% of the M⍰I/V/L mutations came from buried residues whereas only 41.1% of the R⍰K mutations were in buried residues (Data S2). For example, 7 out of 9 R⍰K mutations identified in *rps2* were surface residues while only 1 out of 3 M⍰I/V/L mutations was on the protein surface (Fig. 4F).

Second, controlled *in silico* mutation assays on reconstructed ancestral sequences allowed us to tease apart the complex interplay of protein subunit, mutation type, and residue position quantitatively using Linear Mixed-Effects Models (LMM; see Methods) (Fig. 4C). The positions of these artificial mutations were referenced from true mutations observed in empirical data (Data S2). The best-fitting model suggested that M⍰I/V/L and R⍰K mutations generally destabilized protein structure (intercept ΔΔG = 0.838, LMM p-value = 4.1e⁻⁵). The ΔΔG was significantly influenced by mutation type (i.e., M⍰I/V/L or R⍰K; LMM p-value = 6.0e^-3^) and subunit identity (i.e., small or large subunit; LMM p-value = 0.017) but not residue position (i.e., surface or buried; LMM p-value = 0.32). In alignment with the findings above, we found that M⍰I/V/L on average had a stronger negative impact on protein stability (median ΔΔG = 0.542) than R⍰K mutations (median ΔΔG = 0.150; Fig. S3B). In one extreme case, the M54I substitution within the *rpl16* protein of the 50-kb inversion clade of Papilionoideae yielded a net loss of 4 local hydrogen bonds (Fig. S4) and thus a destabilized *rpl16* protein (ΔΔG = 0.126). Similar hydrogen bond disruptions were also found in R⍰K mutations. For example, in *rps2,* the R30K and R90K mutations broke the hydrogen bond with nearby amino acid residues (Fig. 4H-I) and the R25K mutation led to the loss of the hydrogen bond with the U798 nucleotide in 16S rRNA (Fig. 4J). Finally, unlike the results above by comparing extant proteins, the proportion of buried versus surface residue position did not differ significantly between M⍰I/V/L and R⍰K mutations or explain the overall ΔΔG (p = 0.32). Thus, additional factors such as the innate biophysical properties of M versus R may drive the differentiated impact on structural stability. In addition, we also observed a difference between the small versus large subunits of PRP (LMM p-value = 0.017). Even within R⍰K mutations, mutations in the large subunit had a significantly greater impact on ΔΔG (median ΔΔG = 0.483; Wilcoxon test FDR adjusted p-value = 5.70e^-6^) compared to the small subunit (median ΔΔG = −0.157; Fig. 4C).

### The correlation between GC content, synthesis cost, and amino acid composition

To further understand the potential causes of amino acid composition shift, we first tested whether the observed trend reflected the direction of the background mutational bias. Among the four plant lineages, only Cymbarieae and Geraniaceae demonstrated significant increases in genome and coding sequence GC content compared to close relatives (Wilcoxon test FDR adjusted p-value < 0.026; Table S1). We used LMM to test for the correlation between GC content and the top four amino acids with GC-rich codons: alanine, proline, glycine, and arginine (R). For CpPRP, none of these four amino acids showed significant correlation with the GC content (Fig. S5; LMM p-value >0.089). For non-ribosomal plastid proteins, however, alanine, proline, and glycine showed significant positive correlation with the overall coding sequence GC content (Fig. S5; LMM p-value < 0.011) and the GC content explained 11.6% to 48.9% of the amino acid frequency variation based on the marginal *r*^2^ (Fig. S5). No correlation between the frequency of R and GC content was found in any tests. Instead, the reduced use of GC-rich amino acids such as R in CpPRP ran counter to the trend of increased genomic GC content in Cymbarieae and Geraniaceae.

We then evaluated whether substitutions associated with relaxed selection could lead to significantly reduced synthesis cost compared to the background. We estimated the changes in synthesis cost along each branch based on inferred amino acid substitutions and the protein cost estimated by Chen and Nielsen (2022) (see Methods). When treating all amino acids as equal, no difference in the net synthesis cost change was detected in the foreground (LMM p-value = 0.78). However, when we separated conservative substitutions between biophysically similar amino acids from non-conserved ones based on the BLOSUM 62 matrix (Henikoff and Henikoff 1992), we found that across the tree, conservative substitutions including R⍰K were significantly biased towards the direction that reduced the synthesis cost, while non-conserved substitutions were biased towards cost increase (LMM p-value = 2.28e^-7^; Fig. 5). Importantly, the effects of selection in the foreground branches were only significant in the conservative substitution category (LMM coefficient = −5.94; p-value for the interaction between selection and substitution type = 0.035). Branches with relaxed selection were also characterized by an enrichment of conservative substitutions and a reduction in non-conserved substitutions compared with the background (LMM coefficient = 3.60 in conservative and −2.33 in non-conserved substitutions), which aligned well with the observed dominance of R⍰K and M⍰I/V/L substitutions in the foreground lineages.

**Figure 5.**
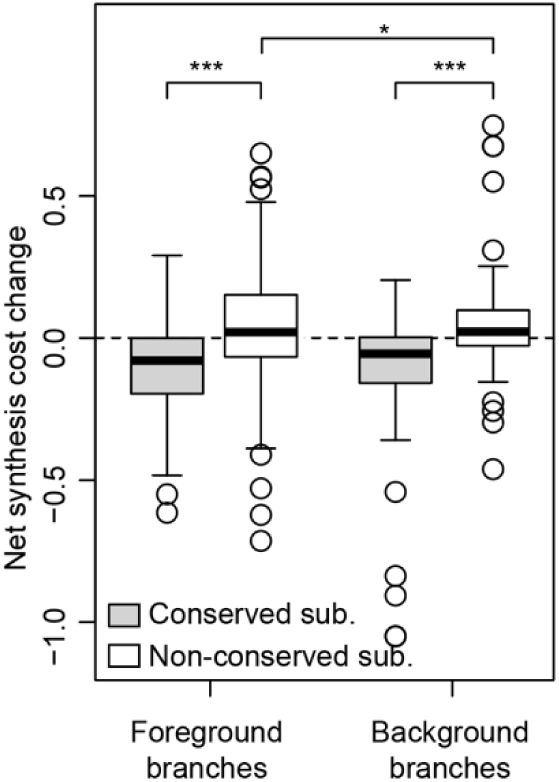
Reduced synthesis cost of amino acids in conserved substitutions but not non-conserved substitutions in plastid ribosome proteins. Net synthesis cost changes were calculated by summing up the per-amino-acid cost change (Chen and Nielsen 2022) for all substitutions along that branch. The horizontal dotted line is a reference for zero changes in synthesis cost. The box plots showed the net synthesis cost changes caused by conserved substitutions (BLOSUM 62 scores >0; gray box) and non-conserved substitutions (BLOSUM 62 scores <0; white box) in foreground and background branches, respectively. Here, the foreground included internal and terminal branches in four plant lineages with relaxed selection in their plastid ribosome proteins. The Linear Mixed-Effects Model (LMM) supported significant difference in synthesis cost between conserved and non-conserved substitutions (p-value = 2.28e^-7^) and significant interaction between the foreground branch type and non-conserved substitutions (p-value = 0.035).

## DISCUSSION

### Plastid genome instability likely triggers coordinated cytonuclear rate shifts in ribosome proteins

Concerted cytonuclear rate shifts in PRP have been reported in multiple plant lineages, including Geraniaceae, Silene, and the 50-kb inversion clade of Papilionoideae (Sloan et al. 2014; Weng et al. 2016; Tressel et al. 2025). This inspired us to follow up on our previous work, where elevated ω values of PRP were found in a single species Monochasma sheareri in Orobanchaceae, to comprehensively investigate PRP evolution within this family (Cai et al. 2026). Here, we demonstrated that the elevated rates evolved in the common ancestor of tribe Cymbarieae. After correcting for RNA editing, significantly higher d_N_, but not d_S_, contributed to the increase in ω in the CpPRP genes of Cymbarieae. This plastid gene rate shift was accompanied by increased d_N_ and ω in the nuclear counterparts of PRP (Fig. 1). The elevated substitution rates in Cymbarieae are thus caused by relaxed selection acting on both plastid-and nuclear-encoded PRP, rather than by changes in background mutation rate. The range of ω and the *k* values (*k* < 1) from the RELAX analyses (Tables S3, S6) further corroborated the relaxed selection scenario. We occasionally obtained high values of ω (> 1) and *k* (> 1) in individual genes such as *rps18* (Tables S5-6), which could be an artifact of short gene sequences. When we concatenated genes by functional groups, both the small and large ribosomal subunits had *k*< 1 (Table S6), suggesting relaxed selection. The exceptionally low values of *k* (1.04e^-7^) from the small subunit suggested more pronounced changes in this complex compared to the large subunit (*k* = 0.475). This could also result from the higher robustness of the small subunit to deleterious mutations as evidenced by our *in silico* mutation assays (Fig. 4C; Fig. S3A). In the small subunit gene *rps2,* introduction of up to 31 substitutions (13.1% of total length) in *Silene noctiflora* still yielded a highly similar structure compared to the ancestral protein (Fig. 4G).

Multiple mechanisms have been proposed to explain such concerted rate shifts, including the relaxed-selection hypothesis as we concluded above (Weng et al. 2016; Tressel et al. 2025). Alternatively, compensatory changes in the nuclear genome in response to rapid plastid evolution have also been proposed (Sloan 2015), but have received less support compared to its application in mito-nuclear interactions (Osada and Akashi 2012). One remaining puzzle regarding the relaxed selection hypothesis is the molecular and ecological causes for such rate shifts. Previous studies have raised a possible link between plastid genome instability and relaxed selection on plastid translation (Schwarz et al. 2017; Tressel et al. 2025). Indeed, all four plant lineages share higher rates of genomic rearrangement, sometimes across both plastid and mitochondrial organelles. Weng et al. (2016) proposed that abundant indels in the 16S rRNA of Geraniaceae plastid genome—the scaffold of the ribosome—may cause substantial structural changes in its protein subunits, thereby leading to relaxed selection on PRP. The 50-kb inversion Papilionoideae similarly have abundant indels and plastid genome size variation likely originating from the genomic shock of the ancestral large-scale inversion (Doyle et al. 1996; Cardoso et al. 2012; Schwarz et al. 2017; Choi et al. 2022). For *Silene* (Caryophyllaceae), species such as *S. conica* and *S. noctiflora* show dramatically shuffled and expanded organelle genomes accompanied by exceptionally elevated gene substitution rates, which are most substantial in mitochondria but also significant in plastids (Sloan et al. 2014). Finally, Cymbarieae is a hemiparasitic lineage with reduced photosynthetic activity (e.g., pseudogenization of photosynthetic complexes in Fig. 1) and frequent local rearrangements of the plastid genome (Ma et al. 2024). It is possible that the erosion of photosynthetic capacity in Cymbarieae relaxed selective constraints on the plastid translational machinery, permitting the accumulation of deleterious mutations in its PRP. We predict that more cases of relaxed selection in PRP will be discovered in plants. Comparative analysis of these independently evolved cases will reveal whether there is a common mechanism, such as plastid instability, behind the relaxed selection on this otherwise highly conserved protein complex.

### Convergent shift of amino acid composition contributes to destabilized plastid ribosome

Across all four angiosperm lineages, shifts in selective pressure are accompanied by consistent changes in amino acid usage. Most significantly, we found increased use of K and reduced use of M driven by high rates of R⍰K and M⍰I/V/L substitutions. Such shifts cannot be explained by background mutational bias or GC content as proposed in other groups (Knight et al. 2001; Palacios and Wernegreen 2002; Horton and Taylor 2023). Thus, the shifts in amino acid composition in PRP were caused by factors beyond the background mutation bias.

A closer look at dominant substitutions like R⍰K revealed that ancestral amino acids were replaced by biophysically similar alternatives with lower molecular weight and shorter side chains (Fig. 3). For example, R and K share similar positive charge, polarity, and molecular volume, whereas M, I, V, and L all have hydrophobic side chains (Grantham 1974). Their positive substitution scores in the BLOSUM62 matrix further indicate that these amino acids are readily exchangeable even in conserved proteins (Henikoff and Henikoff 1992). In proteins like *rps2,* we found multiple parallel R⍰K and M⍰I/V/L substitutions at sites 14, 25, 30, 90, 128, and 135 (Fig. 4E), suggesting high exchangeability at these positions. However, the use of alternative amino acids with lower molecular weight still brings about subtle, but significantly negative impact on the structure and potentially function of proteins (Fig. 4). Methionine (M), along with I, V, and L, are common structural residues that constitute the hydrophobic core of the protein, where purifying selection is typically strongest compared to surface residues (Franzosa and Xia 2009; Tóth-Petróczy and Tawfik 2011). Methionine confers additional stabilization compared to I/V/L because its unbranched and deformable side chain can fill irregular internal cavities and thus optimize protein packing (Lim et al. 2019), which is especially beneficial for the tightly packed PRP (Fig. 4D). In addition, the polarizable sulfur atom in M may engage with diverse nearby aromatic residues to further enhance structural stability (Aledo 2019). In contrast, I, V, and L are rigid β-branched residues that may induce steric conflicts during packing and thus may not be favored when structural integrity is under strong selection. Our prediction from ELASPIC2 strongly supported the deleterious effects of M⍰I/V/L substitutions, with 69.1% predicted to destabilize the protein (Fig. 4A). The extreme case of M54I substitution in *rpl16* led to the loss of eight hydrogen bonds (Fig. S4). These lost interactions suggest that M⍰I substitutions can locally disrupt packing and backbone hydrogen-bonding geometry, even when the replacement amino acid is broadly hydrophobic.

On the other hand, near-neutral impacts were predicted for R⍰K substitutions based on ELASPIC2 (Fig. 4A; Fig. S3B). Some R⍰K substitutions, such as R30K and R90K in *rps2* (Fig. 4H-I), were inferred to break the hydrogen bonds with nearby amino acids, but most R⍰K substitutions do not negatively impact the protein structure. This was expected because McShea et al. (2025) proposed that R is superior to K not only because it forms more hydrogen bonds that enhance folded protein stability (e.g., Fig. 4H-I), but also because R has a smaller solvent-accessible surface and more protein-protein interfaces that shape the surface properties of the protein (Lins et al. 2003; Levy et al. 2012). Indeed, more than half of the R⍰K substitutions in PRP occur at surface residues (Data S2), many of which directly interact with rRNA or nearby proteins. For example, the R25K mutation in *rps2* repeatedly evolved in Geraniaceae, Papilionoideae, and Cymbarieae (Fig. 4J), which broke the hydrogen bonds with the U798 nucleotide of 16S rRNA as well as the A24 residue of *rps2* (Fig. 4J). These surface interactions were, however, not captured by ELASPIC2 or other currently available tools due to the complexity of modeling protein-rRNA interactions in large multiunit complexes. Thus, our near-neutral predictions of R⍰K substitutions using ELASPIC2 should not be interpreted as evidence that these substitutions have negligible functional consequences. Along these lines, we observed twice as much usage of R in PRP compared to other plastid proteins (Fig. S3). This enrichment is likely maintained by selection to preserve proper PRP function and interaction capacity that specifically favors R.

Finally, the lower evolutionary rates observed in the large ribosomal subunit likely reflect its greater proportion of buried residues. Unlike the small subunit, which has a flatter architecture with more solvent-exposed residues, the large subunit forms a tightly packed globular complex (Fig. 4D), making it more structurally constrained and less tolerant of amino acid substitutions (Berman et al. 2000; Ahmed et al. 2016; Ahmed et al. 2017). Overall, the observed rate variation among sites and subunits in PRPs aligns well with the negative correlation between structural constraint and surface exposure (Franzosa and Xia 2009).

### The dynamic trade-off between mutation, structure, and cost

Our approach of tracking the evolutionary fate of individual proteins has allowed us to tease apart the influence of mutation, structural constraint, and synthesis cost in shaping protein evolution. We first ruled out mutational bias as an explanation for amino acid composition shifts: 1) No difference in the background mutation rate (d_S_) was detected in Cymbarieae, Geraniaceae, or the 50-kb inversion clade of Papilionoideae (Weng et al. 2016; Tressel et al. 2025). 2) The increased GC content in Cymbarieae and Geraniaceae ran counter to the decreased use of GC-rich amino acids like R in these lineages. 3) Despite the positive correlation between GC content and GC-rich amino acids in other plastid genes, no such trend was found in PRP (Fig. S5).

Synthesis cost emerged as a plausible alternative drive (Seligmann 2003). In the focal lineages, the ancestral R and M residues were replaced by amino acids with lower molecular weight and potentially lower synthesis cost. For example, R contains a guanidinium group comprising three nitrogen atoms, whereas K terminates in a single amino nitrogen. This makes R more expensive to synthesize in terms of ATP needed based on the modeling results from *Arabidopsis*, especially under nitrogen-starved conditions (Arnold et al. 2015). When measured by the total biomass of synthesis enzymes (i.e., the protein cost), R is four times more expensive than K (Chen and Nielsen 2022). Similarly, M contains sulfur that needs to be incorporated through energetically expensive sulfur assimilation and transfer reactions, making its biosynthesis particularly costly compared to I, V, and L (protein cost = 0.08097 in M versus ≤ 0.00977 in I, V, and L) (Chen and Nielsen 2022). These dominant substitutions are thus biased towards cost-minimization, which is a well-recognized selective force shaping proteomes in archaea, bacteria, and multicellular eukaryotes (Akashi and Gojobori 2002; Seligmann 2003; Swire 2007; Grzymski and Dussaq 2012). Moreover, our LMM model strongly supported cost minimization in all conservative substitutions besides R⍰K and M⍰I/L/V, but this trend was absent in non-conserved substitutions (Fig. 5). Such a contrasting outcome aligns with the theoretical predictions that constraints on protein function outweigh cost minimization (Seligmann 2003). In lineages like Cymbarieae, structural constraints are further lifted to allow for the accumulation of slightly deleterious substitutions that are shaped by cost minimization and ultimately lead to convergent shifts in amino acid composition. Non-conserved substitutions, on the other hand, were likely maintained by compensatory structural mutation or Darwinian selection instead of cost minimization (Osada and Akashi 2012; Sloan 2015). Finally, the absence of amino acid usage change in the nuclear genome likely reflected the different synthesis costs in the nuclear versus plastid environments, because species with extensive organellar genome reconfiguration often show major reorganization of the organellar translational machinery as well (DeTar et al. 2024; Ceriotti et al. 2026; Warren et al. 2026).

The effects of cost-minimization have been demonstrated in bacteria, fungi, animals, and even cancer cells (Akashi and Gojobori 2002; Wagner 2005; Swire 2007; Raiford et al. 2008; Grzymski and Dussaq 2012; Zhang et al. 2018; McShea et al. 2025; Kasalo et al. 2026), but studies in plants are still sparse. Unlike animals, (most) plants are sessile autotrophs that are capable of synthesizing all 20 amino acids. Thus, the price tag behind amino acid production may be different for plants compared to animals; the latter of which can outsource expensive amino acids (Kasalo et al. 2026). Our case study of PRP provides a new empirical example of how this mass-produced protein complex is synergistically shaped by functional and cost constraints. Whether cost minimization is a pervasive force across plant proteomes or is largely restricted to highly expressed proteins remains an open question. Addressing this will require more realistic and biologically relevant models of amino acid synthesis cost in plants because most current models are derived from unicellular organisms and only consider “cost” from the point of chemical energetics (Wagner 2005; Heizer et al. 2006; Raiford et al. 2008). Here, we adopted the protein cost metrics from Chen and Nielsen (2022) for quantification because we believe that they more realistically reflect the nitrogen and carbon investment of protein biosynthesis in plants, which are known to drive their genome size evolution (Kelly 2018; Bales and Hersch-Green 2019; Majda et al. 2021; Hersch-Green et al. 2025). However, the yeast-based protein cost still represents an overly simplified measurement, which does not take into account plant-specific reactions of amino acid biosynthesis and the acquisition of raw substrates by plants with diverse ecological strategies. It is notable that in vertebrates, using yeast-derived chemical energy to measure synthesis cost can lead to contrasting conclusions (i.e., R is more costly than K), even though similar preferences of R over K and I over V were discovered (McShea et al. 2025).

## Conclusion

We established that the hemiparasitic tribe Cymbarieae undergoes concerted cyto-nuclear rate elevation in its plastid ribosomal proteins (PRP). Relaxed selection on PRP has led to convergent shifts in amino acid composition in four distantly related plant lineages including Cymbarieae (Orobanchaceae), *Silene* (Caryophyllaceae), Geraniaceae, and the 50-kb inversion clade of Papilionoideae (Fabaceae). These compositional shifts can be attributed to abundant R⍰K and M⍰I/L/V substitutions, which systematically favor replacement residues that are biosynthetically cheaper and require lower enzymatic biomass to synthesize. Using deep-learning-powered protein structure modeling, we found that these cost-saving molecular adjustments generally reduced protein folding stability and were thus more abundant in surface residues less sensitive to deleterious mutations. Overall, the molecular evolution trend, protein structural prediction, and estimated biosynthesis cost of amino acids demonstrate that biosynthesis cost acts as a pervasive secondary selective driver, actively remodeling highly expressed protein complexes when structural constraints are relaxed. Finally, we highlight the need for biologically realistic cost estimation models for plants to untangle the nuanced interplay between structural constraints, metabolic costs, and mutational biases in driving plant proteome evolution.

## METHODS

### Taxon sampling of Orobanchaceae plastid

We sampled plastid genomes from 21 Orobanchaceae, 2 Paulowniaceae, and 1 Phrymaceae as outgroup (Table S1). Within Orobanchaceae, we select species with diverse lifestyle strategies and represent all major tribes sensu Fischer (2004). This included 4 species from tribe Rehmannieae, 1 from tribe Lindenbergieae, 5 from tribe Cymbarieae, 2 from tribe Orobancheae, 3 from tribe Buchnereae, 1 from genus *Brandisia*, 2 from tribe Rhinantheae, and 3 from tribe Pedicularideae (Table S1). Our primary focus was tribe Cymbarieae, for which we sampled four of the five recognized genera: *Cymbaria* L., *Monochasma* Maxim., *Siphonostegia* Benth, and *Schwalbea* L. (Fig. S1).

We generated plastid genomes from *Monochasma sheareri*, *Alectra sessiliflora* R.Br., and *Melasma physalodes* (L.) Pennell and described these genomes in a companion paper (Cai et al. 2026). All other plastid genomes were downloaded from public NCBI GenBank records (Table S1). Because some parasitic plant taxa have lost, pseudogenized, or partially highly divergent plastid genes due to the evolution of parasitism, not every species contributed a sequence for every gene. We recorded such absences (Tables S4-5) and removed unalignable divergent sequence segments from subsequent rate analyses.

### Masking RNA editing sites for molecular rate analysis

Prior to the molecular evolution analysis, we bioinformatically predicted C-to-U RNA editing sites in plastid genes using the web-based PREPACT v3 (http://www.prepact.de/prepact-main.php)(Lenz et al. 2018). Based on 27 reference species from green algae to angiosperms, C-to-U RNA editing sites were predicted for each species using the default BLAST e-value threshold of 1e^-5^ in PREPACT (Table S4). We then used a custom Python script to process the html output from PREPACT to filter for RNA editing sites that were supported in >60% of the reference species (all scripts available in the GitHub repository: https://github.com/lmcai/Cymbarieae_plastid_ribosomal_proteins). These filtered RNA editing sites were masked in the alignment prior to the downstream inference of substitution rates to prevent overestimation of nonsynonymous substitutions. We did not use an RNA-mapping approach to annotate editing sites because of the limited availability of RNA sequencing data in Orobanchaceae, especially in the focal tribe Cymbarieae.

### Sequence alignments and phylogeny

Codon alignments of the masked plastid sequences in Orobanchaceae were generated by MACSE v2.07 (Ranwez et al. 2018), which can accommodate frameshifts and premature stop codons. The resulting alignments were examined manually in AliView v1.30 (Larsson 2014), where we ensured correct reading frames and carefully screened for potential pseudogenes, premature stop codons, and frameshifts. Based on the resulting codon alignments, a maximum likelihood phylogeny was constructed for each gene by IQ-TREE v3.0.1 with the best substitution model determined by the built-in ModelFinder (Kalyaanamoorthy et al. 2017; Wong et al. 2025). The branch support was evaluated by performing 1000 ultrafast bootstrap (Hoang et al. 2018). To infer a species tree for illustrative purposes (Fig. 1A), we concatenated all plastid genes into a super matrix and inferred a maximum likelihood tree using the same IQ-TREE parameter settings described above.

### Orobanchaceae transcriptome assembly

To investigate nuclear-encoded PRP, we obtained published genomes and transcriptomes from 22 Orobanchaceae species that aligned with the genus level sampling of the plastid dataset (Table S2). Among them, transcriptome assembly was performed for 10 species where assembled transcriptomes were not available. RNA sequencing reads were downloaded from the GenBank Sequence Read Archive and filtered using Fastp v.0.20.1 (Chen et al. 2018). Transcriptome was assembled in Trinity v2.15.2 with the -trimmomatic flag to further remove adapters and low-quality reads (Grabherr et al. 2011). Open reading frames and coding sequences were then predicted by TransDecoder v5.7.1 (https://github.com/TransDecoder/TransDecoder) and only the longest reading frame was retained. The quality and completeness of all nuclear genomes and transcriptomes were assessed using the BUSCO v6 land plant database embryophyta_odb12 (Simão et al. 2015) and reported in Table S2.

### Nuclear-encoded plastid-targeted genes in Orobanchaceae

Three sets of nuclear genes were included: nuclear-encoded plastid-targeted ribosomal protein genes (NuPRP), nuclear-encoded cytosol-targeted ribosomal genes (NuCyRP), and other nuclear-encoded plastid-targeted genes that are not involved in ribosomes (NuCpOT). Their homologs from *Arabidopsis thaliana* (L.) Heynh. for each set were identified in the UniProt database (www.uniprot.org; Table S3) and largely followed Tressel et al. (2025) and Weng et al. (2016). Candidate orthologs in Orobanchaceae were obtained by BLASTx search with an e-value threshold of 1e^-20^. The DNA sequences of all BLASTx hits were aligned into codon positions using MACSE and were used to generate maximum likelihood phylogenies in IQ-TREE following the same steps described above. These phylogenies were inspected manually to confirm orthology and remove any non-orthologous sequences.

To prepare nuclear-encoded plastid-targeted genes for rate analyses, we identified the cleavage site of N-terminal peptides using the TargetP 2.0 web server (https://services.healthtech.dtu.dk/services/TargetP-2.0/)(Almagro Armenteros et al. 2019). After removing transit peptides from the codon alignments, we realigned the cleaned sequences using MACSE and generated a final gene tree for downstream analyses.

### Substitution rate estimation in CODEML

The pairwise model of CODEML in PAML v4.10 (Yang 2007) was used to estimate d_N_, d_S_, and their ratio ω per species per gene. In an explorative analysis, we tested the performance of the free-ratio mode to infer individual rates for each branch as implemented in Tressel et al. (2025) and Weng et al. (2016). However, due to the conservative nature of plastid genes, the parameter-rich free-ratio mode frequently generated exceptionally low d_S_ values for internal branches, leading to unrealistic ω estimates (>99) across multiple branches of the phylogeny. We thus chose to use the pairwise mode for rate estimation.

For CODEML analyses, we used plastid alignments with masked RNA editing sites and nuclear alignments without N-terminal peptides. Five gene sets were used: NuPRP, NuCyRP, NuCpOT, CpPRP, and CpPS. Each sequence was aligned to the Phyramaceae outgroup and CODEML analyses were performed under the pairwise mode (runmode = –2) with the F3X4 codon frequency model. To test for rate differences between Cymbarieae and other Orobanchaceae for each function group (e.g., NuPRP), we applied Wilcoxon rank sum tests and adjusted the resulting p-values using false discovery rate correction (FDR).

### Testing relaxed selection

To test for relaxed selection in Cymbarieae, we applied the RELAX method from HyPhy v.4.5 to both individual plastid and nuclear genes as well as concatenated sequences grouped by functional classes (Tables S3 and S6) (Wertheim et al. 2015; Kosakovsky Pond et al. 2020). RELAX compares the ω value in the foreground branches (i.e., Cymbarieae) to the background branches in a reference species tree. If the foreground branches are under relaxed selection, their ω values are expected to converge towards the neutral value 1 across all rate categories and vice versa for intensified selection (Wertheim et al. 2015). We trimmed the tips of the species tree inferred above to align with the taxon sampling of each gene. We selected the internal and terminal branches within Cymbarieae as foreground and tested whether they showed significantly relaxed or intensified selection compared to the background.

### Testing convergent shift in amino acid composition

To characterize convergence in amino acid composition shift in lineages experiencing relaxed selection, we downloaded the sequences of plastid-and nuclear-encoded PRP from Tressel et al. (2025), Weng et al. (2016), and Sloan et al. (2014). This included 40 species from the 50-kb Inversion clade of Papilionoideae (Fabaceae), 25 species from Geraniaceae, 2 species from the fast-evolving *Silene* (Caryophyllaceae), along with outgroups with slower-evolving PRP within each of the three studies (Table S7). These coding sequences were merged with our Orobanchaceae dataset and alignments were generated using MACSE, which resulted in one codon alignment and one amino acid alignment per gene. The maximum likelihood species tree was inferred from the concatenated plastid coding sequence alignments using IQ-TREE following the same steps described above (Fig. S6).

We reconstructed ancestral protein sequences and summarized the changes in amino acid composition over time. To do this, we applied the empirical Bayesian method implemented in IQTREE to infer the ancestral sequences for plastid-and nuclear-encoded PRP genes. The species tree inferred above (Fig. S6) was used as the reference phylogeny for sequence reconstruction. The resulting ‘.state’ files from IQTREE were processed in a custom R script AA_composition_summary.R (available on GitHub) to filter for sites experiencing substitution in at least one descending lineage within the focal group (e.g., Cymbarieae). This allowed us to visualize the overall shift in amino acid composition accompanying relaxed selection. Statistical differences in the usage of the 20 amino acids between the focal lineage and their corresponding outgroups were evaluated using the Wilcoxon rank-sum test, and p-values were adjusted using FDR.

To further characterize the process of amino acid substitution leading to shifts of overall composition bias, we used a custom Python script AA_flux.py (available on GitHub) to count the number of amino acid substitutions for each internal and terminal branch of the focal lineages based on reconstructed ancestral sequences. We then summed up all substitutions to calculate the total flux rate between any two pairs of amino acids within the focal lineage. The positions of each mutation type were also recorded to inform the *in silico* mutation analysis of protein folding stability below.

### Mutation analysis of protein folding stability

To test how individual amino acid mutations impact the protein folding stability, we performed two sets of complementary analyses to examine the structural impact of amino acid substitutions accompanying relaxed selection. We used the deep-learning-based ELASPIC2 (Strokach et al. 2021) to predict how individual mutations might impact the free energy of protein folding (ΔΔG). A positive ΔΔG value suggests a destabilized structure compared to the reference and vice versa.

First, as a proof of concept, we compared the ΔΔG between species with and without relaxed selection in each of the four lineages. Within each lineage, ΔΔG values were computed from all possible pairwise combinations between a focal species with relaxed selection and a reference species without relaxed selection. Each pairwise comparison contained multiple mutations and the resulting ΔΔG value reflected the additive effects of individual mutations as well as their epistatic interactions.

Secondly, we reconstructed ancestral sequences and introduced artificial mutations to perform a more controlled experiment. We specifically focused on two types of mutations, R⍰K and M⍰I/V/L, which showed consistent trends across all four plant lineages. We used the ancestral sequence from the above section (‘Testing convergent shift in amino acid composition’) and added one R⍰K or M⍰I/V/L mutation each time based on mutations observed in empirical sequences. We manually inspected the sequence alignment to prioritize mutations located in conserved domains and exclude those nested within stretches of multiple mutations or indels, because the latter mutations were likely results of larger-scale structural changes. As a result, each pairwise comparison contained only one mutation to the reference and the resulting ΔΔG value reflected the structural impact of individual mutations.

To assess the thermodynamic stability effects of mutations, we employed ELASPIC2 (Strokach et al. 2021) to predict the change in ΔΔG (measured in kcal/mol) by introducing mutation(s) to a reference protein structure. ELASPIC2 combines pre-trained transformer-based language models with graph convolutional neural networks to generate accurate predictions of mutational effects on protein stability (Strokach et al. 2021). To run ELASPIC2 on sequences with introduced mutation(s), we generated the 3D protein structures using ColabFold (Mirdita et al. 2022) for each reference sequence and manually validated them. The resulting protein structure and identity of the mutation(s) were subsequently used to predict ΔΔG based on structural predictions from ProtBert (Elnaggar et al. 2022) and ProteinSolver (Strokach et al. 2020) in ELASPIC2.

### Structural annotation using the spinach cryo-EM structures

To interpret the ΔΔG predictions in the context of residue 3D positions, we annotated the structural features of each amino acid residue based on the high-resolution cryo-EM structures of PRP from spinach (*Spinacia oleracea* L.). The small (PDB ID: 5X8R) and large subunits (PDB ID: 5H1S) of spinach PRP were downloaded from the RCSB Protein Data Bank (Berman et al. 2000; Ahmed et al. 2016; Ahmed et al. 2017)(https://www.rcsb.org/). We classified each residue in the spinach PRP as buried or surface based on its solvent accessibility estimated by FreeSASA (Mitternacht 2016). FreeSASA applied the Lee-Richards algorithm to calculate the relative solvent accessibility of each residue by normalizing its total accessible surface area against the theoretical maximum values. Residues with relative solvent accessibility larger than 0.25 were classified as "surface" and otherwise as "buried". Finally, we transferred the spinach PRP residue annotation to the homologous sites within each reference sequence. Here, homology was established by sequence alignment using MAFFT as described above. Regions showing substantial divergence from the spinach were discarded (e.g., *rpl23* from spinach was not alignable to any ingroup).

### Quantifying sources of variation in protein folding stability using Linear Mixed-Effects Models

We used the Linear Mixed-Effects Model (LMM) to test the significance of protein subunit type (small versus large), mutation type (M⍰I/V/L versus R⍰K), and residue position (buried versus surface) in influencing the protein folding stability ΔΔG when M⍰I/V/L or R⍰K were introduced to ancestrally reconstructed proteins. We fitted an LMM using the lmer function from the R package lme4 (Bates et al. 2015). The response variable was ΔΔG estimated for each introduced mutation, and fixed effects included protein subunit type, mutation type, and residue position. Clade was included as a random intercept to account for the non-independence of observations within the four major plant lineages. Significance of fixed effects was assessed using Satterthwaite’s approximation for degrees of freedom as implemented in the lmerTest package (Kuznetsova et al. 2017). All R code used for LMM analysis hereafter was available on GitHub.

### Correlation of GC content and amino acid composition shift

To examine whether the baseline mutational bias, reflected in GC content, may explain shifts in amino acid composition post-relaxed-selection, we applied LMM to test for correlation between selected amino acids and GC content. We especially examine the frequency of the four most GC-biased amino acids—alanine (83.3% codon GC content), proline (83.3%), glycine (83.3%), arginine (72.2%)—to genomic and coding sequence GC content. The response variable was amino acid frequency, and fixed effects included GC content. Clade was included as a random intercept to account for the non-independence of observations within the four major plant lineages (i.e., Orobanchaceae, Fabaceae, Geraniaceae, and *Silene*). We ran the test using the lmer function and tested the correlation using the amino acid frequency from CpRP and all other plastid proteins, respectively.

### Correlation between synthesis cost, amino acid substitution type, and relaxed selection

To demonstrate that reduced synthesis cost was preferred under relaxed selection, we applied LMM to examine the correlation between net synthesis cost change and relaxed selection. The expectation under this hypothesis is that branches with relaxed selection should have reduced net synthesis cost or more cost-reducing substitutions.

Based on the ancestral sequence from the above section (‘Testing convergent shift in amino acid composition’), we mapped individual substitutions to each internal and terminal branch. Then, we calculated the normalized synthesis cost flux per substitution for each branch based on the protein cost of amino acid synthesis estimated by Chen and Nielsen (2022):

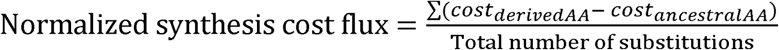

Initial LMM analysis revealed an insignificant difference in the normalized synthesis cost flux in foreground versus background branches (LMM p-value = 0.78). However, we noticed that the substitution type might have a major impact. We then classified the 380 pairwise substitutions into two types: 1) conserved substitutions between amino acid pairs with a positive BLOSUM 62 score; 2) the non-conserved substitutions between amino acid pairs with a negative BLOSUM 62 score (Henikoff and Henikoff 1992). To evaluate whether the synthesis cost differed based on selection and substitution types, we fitted an LMM using the lmer function. The response variable was normalized synthesis cost flux per substitution, and fixed effects included selection type, substitution type, and their interaction. Clade was included as a random intercept to account for the non-independence of observations within the four major plant lineages. Branch length was incorporated as an observation-level weight, such that estimates were weighted according to the evolutionary time represented by each branch. Significance of fixed effects was assessed using Satterthwaite’s approximation for degrees of freedom as implemented in the lmerTest package (Kuznetsova et al. 2017).

## Acknowledgement

The authors thank Stuart McDaniel and the EvolDoers discussion group for feedback on the initial version of the study and the computing cluster HiperGator at the University of Florida for the computational resources to complete the study. Funding for this research is provided by the startup fund from the University of Florida to LC; NIH grant R35 GM154908 to JZ and PS.

