## Supplementary Information for "Synthesis cost is a hidden driver of convergent amino acid composition in plastid ribosomal proteins"

**Supplementary figures**

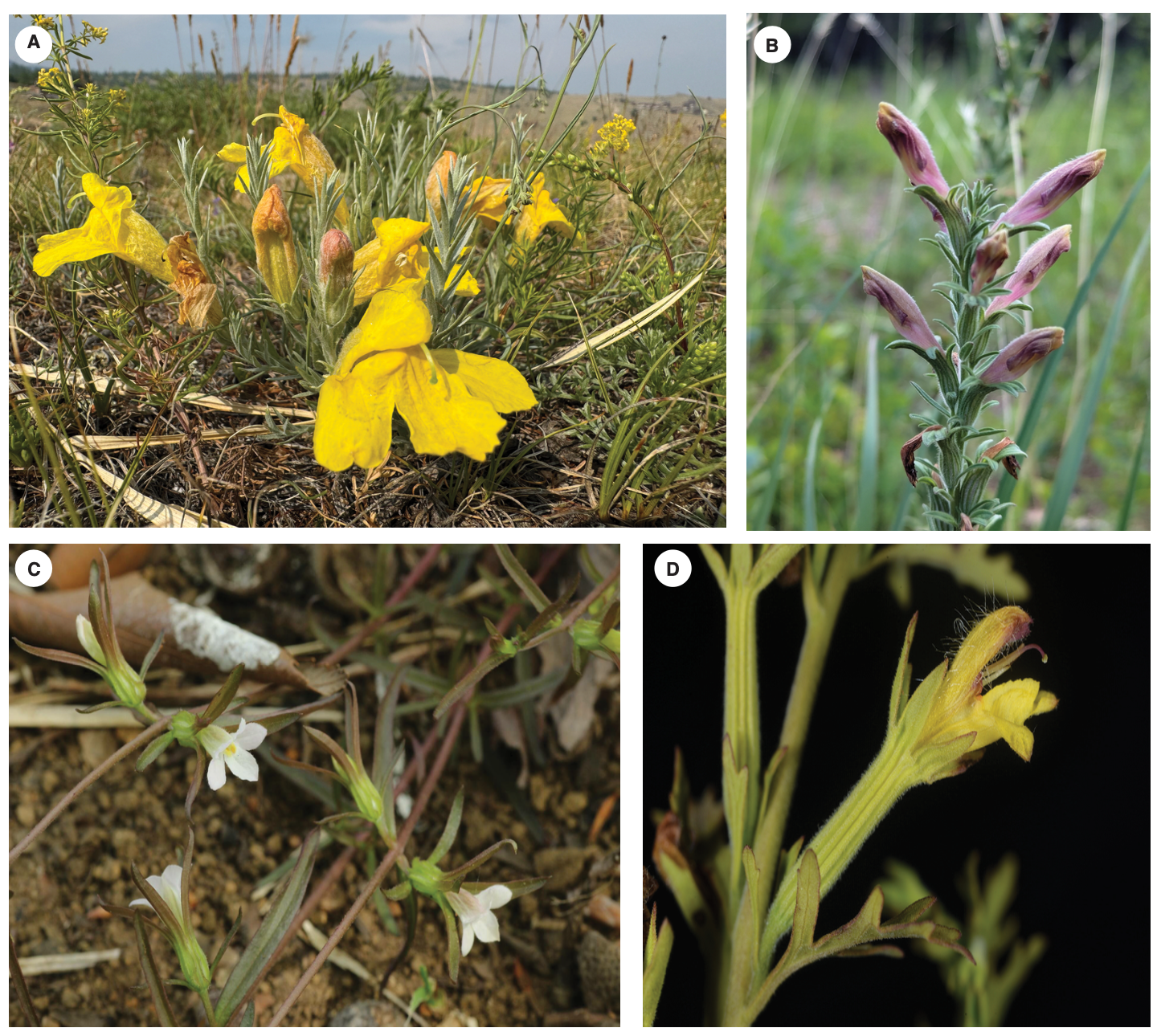

**Figure S1** Representative species of tribe Cymbarieae (Orobanchaceae) illustrating the morphological and ecological diversity across the lineage. (A) *Cymbaria daurica* L., a hemiparasitic herb with bright yellow corollas, occurring in temperate grasslands of northern China (credit: Natalya Kaschuk, CC-BY-NC). (B) *Schwalbea americana* L., a federally endangered hemiparasite with pale yellow to purple-tinged flowers, native to pine savannas and coastal plain grasslands in eastern North America (credit: d0ugh0ck, CC-BY-NC). (C) *Monochasma sheareri* (Maxim.) Franch. ex Maxim., a hemiparasitic species with pink to white tubular flowers, distributed across open forest margins in East Asia (credit: tosakah, CC-BY-NC). (D) *Siphonostegia chinensis* Benth., an annual hemiparasite bearing small yellow corollas, typically inhabiting sandy or disturbed habitats in subtropical east Asia (credit: lecanorchis, CC-BY-NC).

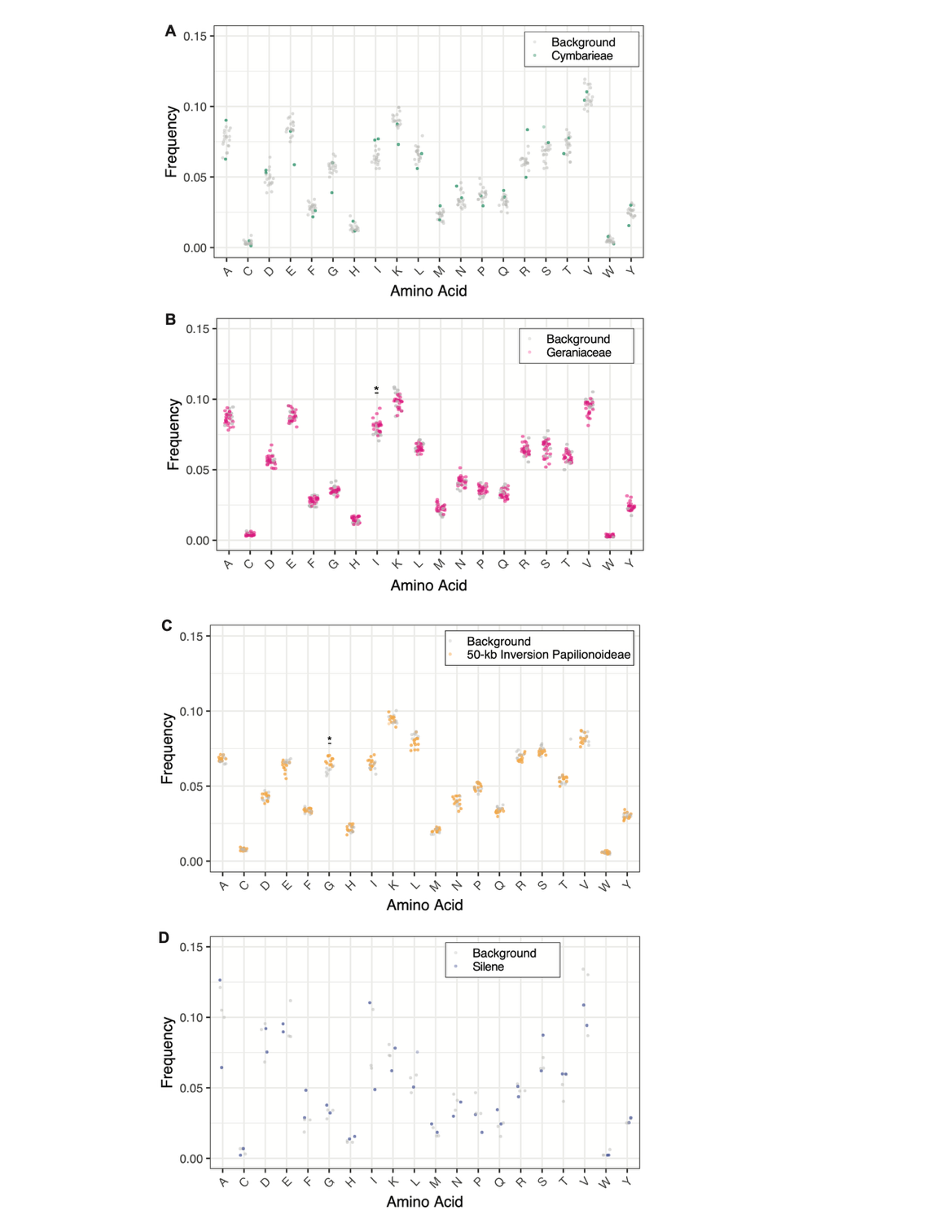

**Figure S2** Amino acid usage of nuclear-encoded ribosomal proteins after relaxed selection. Frequencies of amino acid usage are contrasted between species experiencing relaxed selection (colored dots) and background species (gray dots) in four plant lineages: A. Cymbarieae (Orobanchaceae); B. Geraniaceae; C. 50-kb inversion clade of Papilionoideae (Fabaceae); D. fast-evolving *Silene* (Caryophyllaceae). Asterisks indicate significant differences in amino acid frequency based on Wilcoxon rank-sum tests (* FDR-adjusted p-value < 0.05).

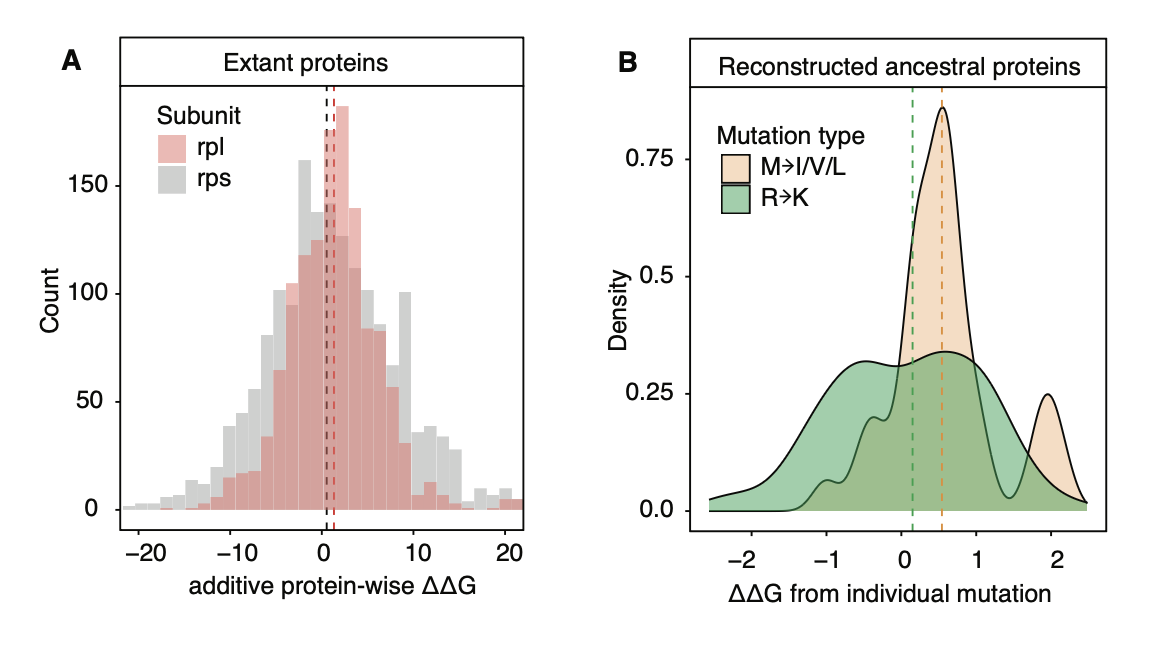

**Figure S3** Impact of M→I/V/L and R→K mutations on plastid ribosome protein structure measured by free energy of protein folding (ΔΔG) in ELASPIC2. Positive ΔΔG suggests destabilized proteins. Dotted lines indicate the median value of ΔΔG for each category. (A) Mutations in the large subunit have stronger destabilizing effects than the small subunit. ΔΔG was inferred from pairwise comparisons of protein structure from extant species with and without relaxed selection. (B) M→I/V/L mutations have stronger negative effects on protein stability compared to R→K mutations. ΔΔG was inferred by introducing individual M→I/V/L or R→K mutations to the reconstructed ancestral protein of each lineage.

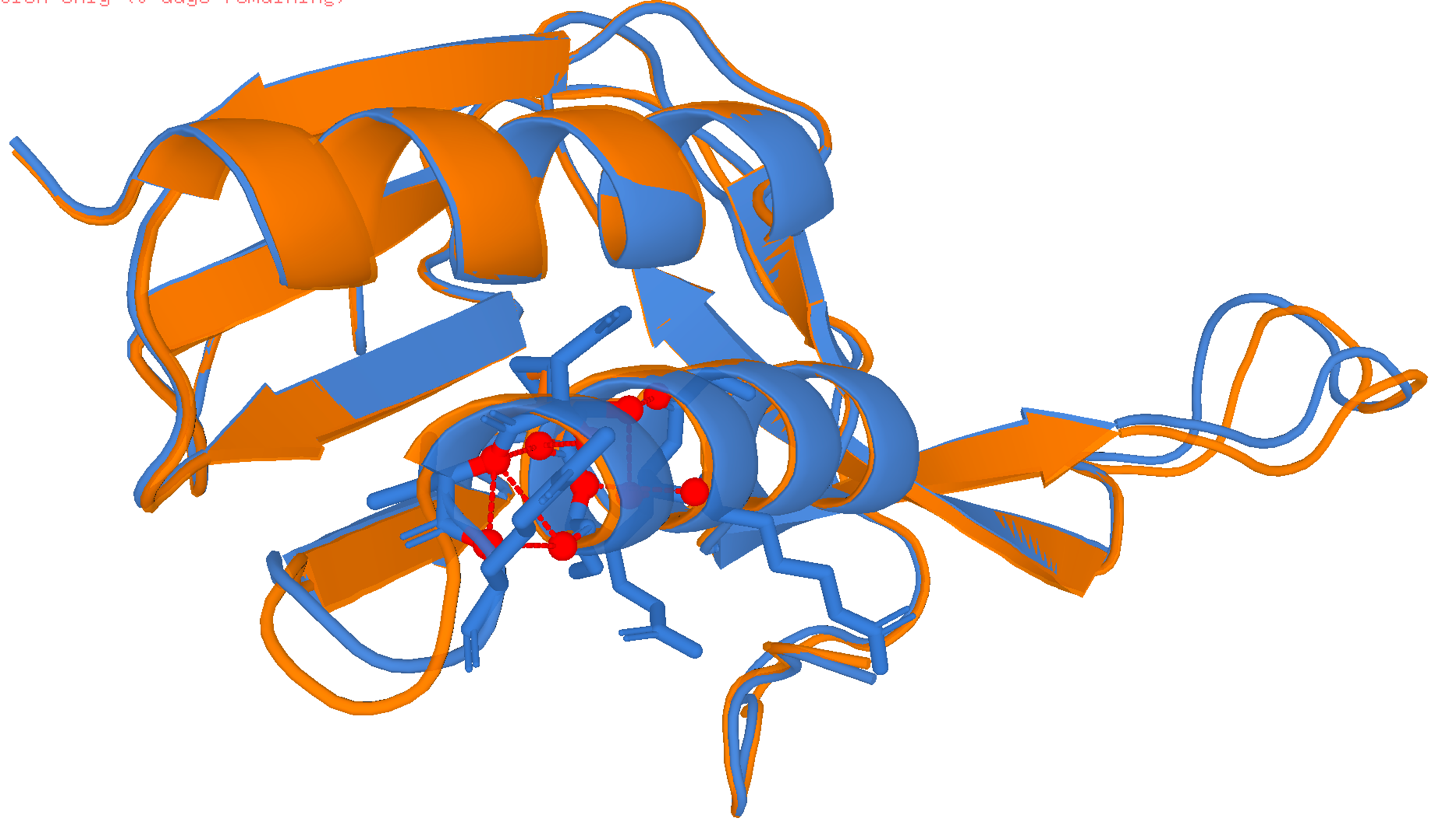

**Figure S4** Structural comparison of ancestral and mutant *rpl16* models around the M54I substitution within the 50-kb inversion clade of Papilionoideae (Fabaceae). The ancestral structure is shown in blue and the mutant structure in orange. Red dashed lines indicate hydrogen bonds present in the ancestral model but absent in the mutant model, restricted to interactions with at least one endpoint within 6 Å of the residue-54 Cα atom. The M54I substitution was associated with eight lost and four gained local hydrogen bonds, corresponding to a net loss of four hydrogen bonds near the mutation site.

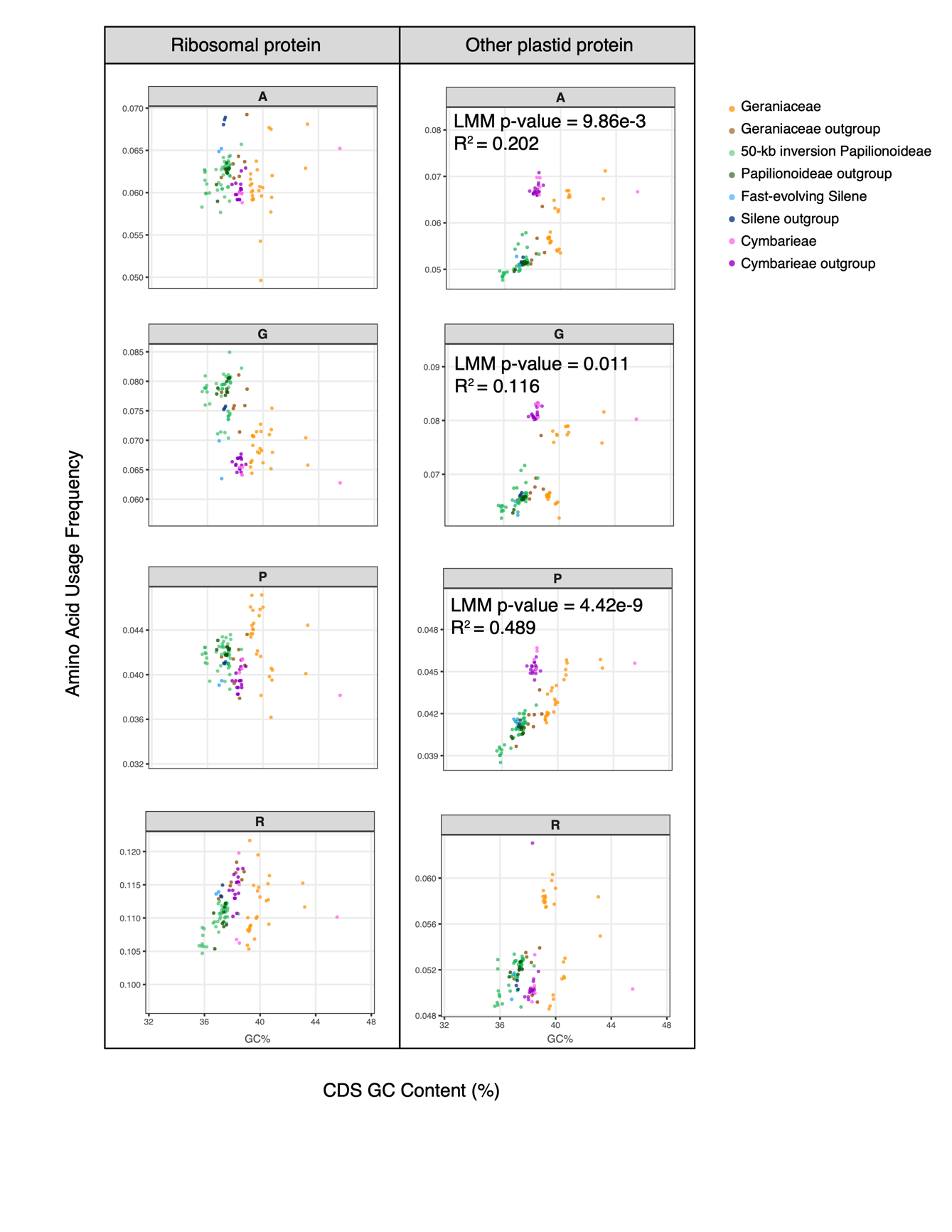

**Figure S5** Correlation between GC content and amino acid composition in plastid genes. Scatter plots of the gene coding sequence GC content (CDS GC%) and amino acid composition (measured in usage frequency) were shown for four amino acids with the most GC-rich codons: alanine (A), glycine (G), proline (P), and arginine (R). Results for plastid-encoded ribosome proteins and all other proteins were plotted separately. For comparisons with significant correlation between CDS GC% and amino acid usage, the p-value and marginal *r*^2^ of variance explained by fixed effects under the Linear Mixed-Effects Model (LMM) were shown in the top left corner.

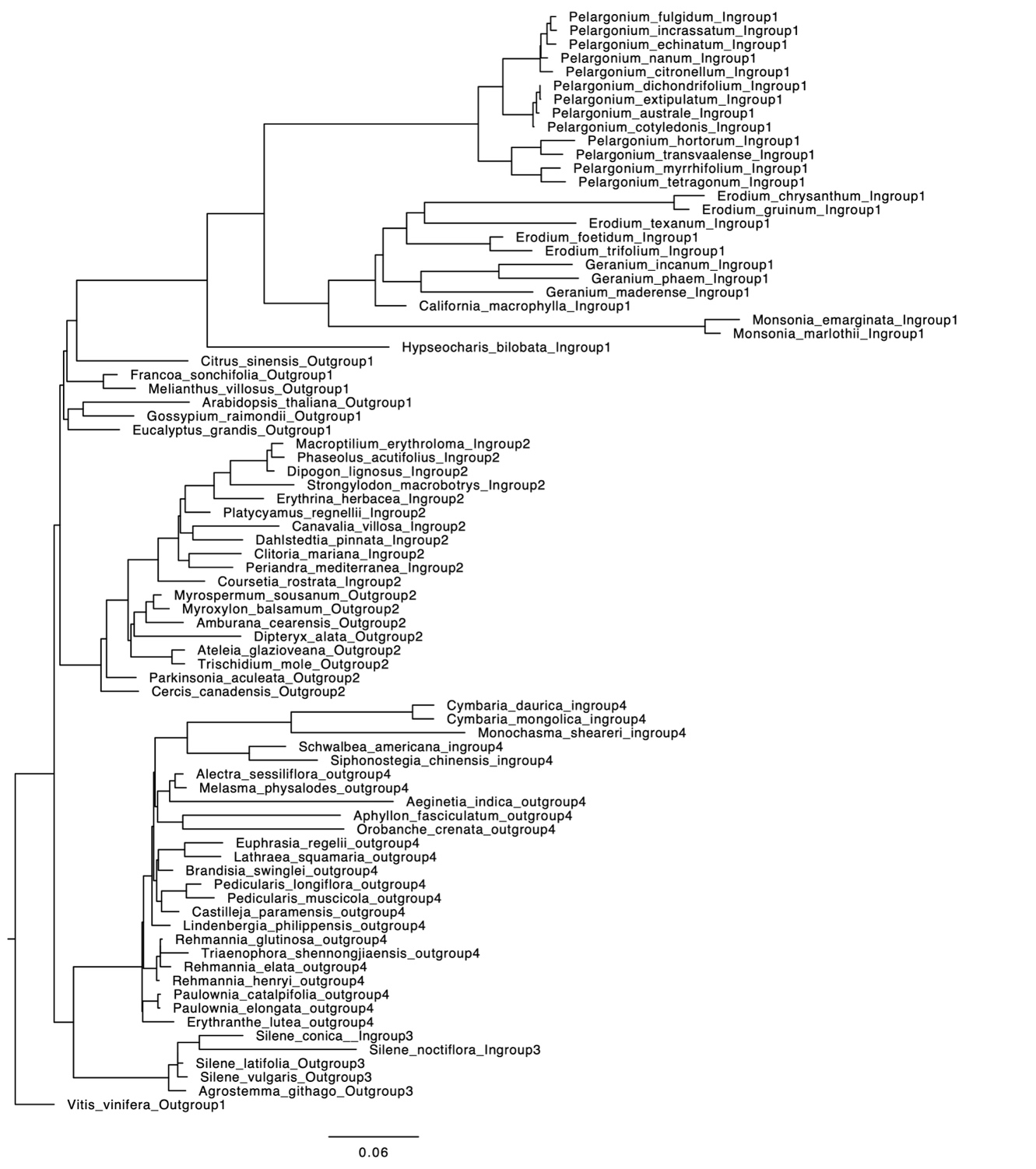

**Figure S6** Maximum likelihood phylogeny of lineages showing relaxed-selected plastid ribosome proteins. The maximum likelihood species tree was inferred from the concatenated plastid coding sequence alignments and the best substitution model determined by ModelFinder in IQ-TREE. ‘Ingroup’ suggests species with relaxed selection and ‘outgroup’ indicates closely related reference species.

**Supplementary tables**

**Table S1** Taxon sampling and data source for plastid genomes in Orobanchaceae and its close relatives.

**Table S2** Taxon sampling, data source, and data quality evaluation for the genomic survey of nuclear-encoded plastid genes in Orobanchaceae. BUSCO assessment of data quality is based on the land plant embryophyta_odb12 database.

**Table S3** Statistical significance of relaxed or intensified selection of nuclear-encoded plastid-targeted ribosomal proteins, nuclear-encoded cytosol-targeted ribosomal proteins, and nuclear-encoded plastid-targeted other proteins in Cymbarieae (Orobanchaceae) using HYPHY. The relaxation/intensification parameter (K) and p-values of test branches (Cymbarieae) are reported for individual genes. Significantly relaxed selection is highlighted in orange and significantly intensified selection is highlighted in blue. The cleavage site of N-terminal peptides was predicted using targetP and removed before the HYPHY analysis.

**Table S4** Number of RNA editing sites per gene predicted by PREPACT for 80 plastid protein coding genes. "N/A" represent loss or psedogenization of the gene.

**Table S5** Synonymous substitution rates (dS), non-synonymous substitution rates (dN), and omega values of plastid genes inferred from pairwise sequence comparison to *Erythranthe lutea* in CODEML.

**Table S6** Statistical significance of relaxed or intensified selection of plastid genes in Cymbarieae (Orobanchaceae) using HYPHY. The relaxation/intensification parameter (K) and p-values of test branches (Cymbarieae) are reported for individual genes. Significantly relaxed selection is highlighted in orange and significantly intensified selection is highlighted in blue.

**Table S7** Taxon sampling and data source for additional taxa with rate shifts in their plastid ribosomal proteins. Taxaon sampling was quoted from Tressel et al (2025) for Papilionoideae, Weng et al (2016) for Geraniaceae, and Sloan et al (2013) for *Silene*.

**Table S8** Evaluation of amino acid usage difference in four plant focal clades and their close relatives based on Wilcoxon rank-sum tests. The raw p-value (p_raw) and FDR-adjusted p-value (p_adj) were reported.

**Table S9** Lack of convergent site substitution in lineages experiencing relaxed selection in plastid ribosome proteins in CSUBST. In both foreground (lineages with relaxed selection) and non-foreground branches, the number of convergent branch combinations with the convergence parameter ω_C_ ≥3 and the number of convergent amino acid substitutions O_C_^N^ ≥3 were included. None of the genes showed significantly enriched convergent branches based on Fisher’s exact test with FDR adjustment (p=1 for all genes).

**Supplementary data**

**Data S1** Sequence alignments of all surveyed plastid and nuclear genes in Orobanchaceae, Geraniaceae, Fabaceae, and Caryophyllaceae. These genes included the plastid-encoded plastid ribosome genes (CpPRP), nuclear-encoded plastid-targeted ribosomal proteins (NuPRP), nuclear-encoded plastid-targeted non-ribosomal proteins (NuCpOT), and nuclear-encoded cytosolic ribosomal proteins (NuCyRP). The RNA editing sites of plastid genes were predicted using PREPACT v3 and masked using a custom Python script. The N-terminal peptides of nuclear-encoded plastid-targeted genes were predicted using TargetP v2.0 and trimmed.

**Data S2** Changes in protein folding stability estimated by ELASPIC2. Protein stability was measured by changes in free energy of protein folding (ΔΔG) using the deep learning powered tool ELASPIC. Pairwise comparisons were performed between 1) observed natural protein pairs (e.g., one Cymbarieae species and a closely related outgroup) and 2) reconstructed ancestral proteins with artificial mutations (e.g., reconstructed ancestral protein of Cymbarieae with introduced R→K mutations).

**Supplementary notes**

**Note S1** **Testing convergent** **amino acid site substitution in PRP using CSUBST**

**Methods**

To test site-level convergence of amino acid substitutions among plants with fast-evolving ribosomal proteins, we analyzed the aligned codon and one amino acid sequences using the program CSUBST v1.4.0 following the method described by (Fukushima and Pollock 2023). These alignments included Cymbarieae (Orobanchaceae), the 50-kb Inversion clade of Papilionoideae (Fabaceae), Geraniaceae, *Silene* (Caryophyllaceae), along with outgroups with slower-evolving ribosomal proteins within each of the four studies. The coding sequences were initially aligned using MACSE v2 (Ranwez et al. 2018), which resulted in one codon alignment and one amino acid alignment per gene. The input phylogeny of CSUBST was the same as the one described in the main text.

Within CSUBST, the rate of amino acid convergence parameter ω_C_ was calculated from the ratio of observed to expected non-synonymous combinatorial substitutions against the ratio of observed to expected synonymous combinatorial substitutions across distinct phylogenetic branches. With the neutral expectation of 1.0, higher ω_C_ values indicate accelerated rates of protein convergence. To prepare the input alignments, we trimmed the amino acid alignment with ClipKIT (Steenwyk et al. 2020) under the gappy mode where sites with more than 70% gaps were removed. We then reverse-translated the trimmed amino acid alignment to the codon alignment with the ‘backtrim’ function of CDSKIT (Fukushima and Pollock 2023). Finally, stop codons and ambiguous codons were replaced with ‘N’ using the ‘mask’ function of CDSKIT to prepare the cleaned codon alignments. To run CSUBST, we set Cymbarieae, the 50-kb inversion clade of Papilionoideae, Geraniaceae, and fast-evolving *Silene* as foreground branches and ran CSUBST under exhaustive combinations of up to ten branches followed by higher-order branch-and-bound search (--max_arity 10). To designate a pairwise branch comparison as convergent, we retained genes and branch combinations with ω_C_ ≥ 3 and the number of convergent amino acid substitutions O_C_^N^ ≥ 3 for any ancestral state transitioning to specific derived states. We tested whether foreground branches were enriched in convergent evolution by applying Fisher’s exact test. Finally, for a few selected genes, the positions of convergent amino acid substitutions were mapped onto their predicted protein structure using the ‘site’ function of CSUBST. All command lines and scripts of data curation and CSUBST were available on GitHub (https://github.com/lmcai/Cymbarieae_plastid_ribosomal_proteins/tree/main/4_convergent_evo_csubst).

**Results and discussion**

**Lack of widespread site-specific convergence in amino acid evolution**

The higher-order branch-and-bound search implemented in CSUBST allowed for detecting any branch combination showing convergent amino acid site substitutions. An enrichment of foreground lineages in the result can serve as evidence for convergence linked with traits unique to the foreground (Fukushima and Pollock 2023). When we restrict the convergent site substitution to the stem group of the foreground lineages only (--fg_stem_only yes) as has been done in (Fukushima and Pollock 2023), no genes showed significant convergence (ω_C_ < 3). When we relaxed the criteria and allowed substitution to occur in both stem and descendant lineages (-fg_stem_only no), six out of 21 (28.6%) CpPRP and 23 out of 29 (79.3%) NuPRP showed significant convergence (ω_C_ ≥ 3 and O_C_^N^ ≥ 3) in at least three foreground or background branches. However, Fisher’s exact test demonstrated that convergence was not enriched in foreground lineages, despite that they accounted for 65.7% of our taxon sampling (FDR adjusted p-value =1 in all tests; Table S9). Furthermore, despite having 110 foreground branches on average, no CpPRP genes show significant convergence among more than 3 branches. For NuPRP genes, 16 genes show convergence among 4 branches and 5 show convergence among 5 branches. No genes show convergence in 6 branches or above out of 110 foreground branches. We thus concluded lack of widespread site-specific convergence in both CpPRP and NuPRP.
